# Cytosolic iron-sulfur protein assembly system identifies clients by a C-terminal tripeptide

**DOI:** 10.1101/2023.05.19.541488

**Authors:** Melissa D. Marquez, Carina Greth, Anastasiya Buzuk, Yaxi Liu, Catharina M. Blinn, Simone Beller, Laura Leiskau, Anthony Hushka, Kassandra Wu, Kübra Nur, Daili J. Netz, Deborah L. Perlstein, Antonio J. Pierik

**Author notes:** Corresponding author. (D.L.P.); (A.J.P.). These authors contributed equally to this work (Melissa D. Marquez, Carina Greth). Present address: MDM: Department of Chemistry Harvard University, Boston, MA; SB: Department of Department of Animal Nutrition and Nutrition Physiology, Justus Liebig University Giessen, Germany; AH: Department of Microbiology, Blavatnik Institute, Harvard Medical School, Boston, MA 02115.

## Abstract

The eukaryotic cytosolic Fe-S protein assembly (CIA) machinery inserts iron-sulfur (Fe-S) clusters into cytosolic and nuclear proteins. In the final maturation step, the Fe-S cluster is transferred to the apo-proteins by the CIA-targeting complex (CTC). However, the molecular recognition determinants of client proteins are unknown. We show that a conserved [LIM]-[DES]-[WF]-COO^-^ tripeptide present at the C-terminus of clients is necessary and sufficient for binding to the CTC *in vitro* and directing Fe-S cluster delivery *in vivo*. Remarkably, fusion of this TCR (target complex recognition) signal enables engineering of cluster maturation on a non-native protein via recruitment of the CIA machinery. Our study significantly advances our understanding of Fe-S protein maturation and paves the way for bioengineering applications.

**One-Sentence Summary:** A C-terminal tripeptide guides eukaryotic iron-sulfur cluster insertion into cytosolic and nuclear proteins.

---

Iron-sulfur (Fe-S) proteins catalyze transformations vital for life-sustaining processes including photosynthesis, respiration, and nitrogen fixation. Although Fe-S cofactors are readily inserted into apo-proteins *in vitro*, their biosynthesis requires dedicated pathways that assemble the metalloclusters and shepherd these cofactors to apo-protein clients (*1*). Cluster biogenesis is essential for iron homeostasis, energy metabolism, genome stability, and other critical cellular processes impacting human health (*2–5*). The diversity of transformations catalyzed by Fe-S enzymes also makes them attractive candidates for bioengineering of novel pathways, however, inefficient cluster delivery to non-native proteins often frustrates pharmaceutical and biofuel applications (table S1) (*6, 7*). Although progress toward understanding the apo-client recognition has been made in some systems (*7–9*), there are no plug-and-play solutions to overcome bottlenecks associated with inefficient metallocofactor maturation.

Eukaryotic Fe-S biogenesis is complex because each subcellular compartment houses its own dedicated machinery (*1*). Cytosolic and nuclear Fe-S protein assembly (CIA) requires several mitochondrial iron-sulfur cluster (ISC) maturation components, including the Nfs1 cysteine desulfurase, and at least 9 additional cytosolic proteins. The ISC machinery synthesizes a molecule containing iron and/or sulfur (XS; Fe-S_int_) (*10, 11*). Upon export, several protein factors use this molecule for [4Fe-4S] cluster assembly on the Cfd1-Nbp35 CIA scaffold (Fig. 1A) (*12, 13*). Nar1, a 2x[4Fe-4S] protein acts as the Fe-S cluster carrier, associates with the CIA targeting complex (CTC, Fig. 1A) and provides the cluster to be inserted into apo-Fe/S targets (*14*). These targets are recognized by the CTC, a complex comprising Met18/MMS19 (yeast/human nomenclature), Cia1/CIAO1, and Cia2/CIAO2 (*15–18*).

**Fig. 1.**
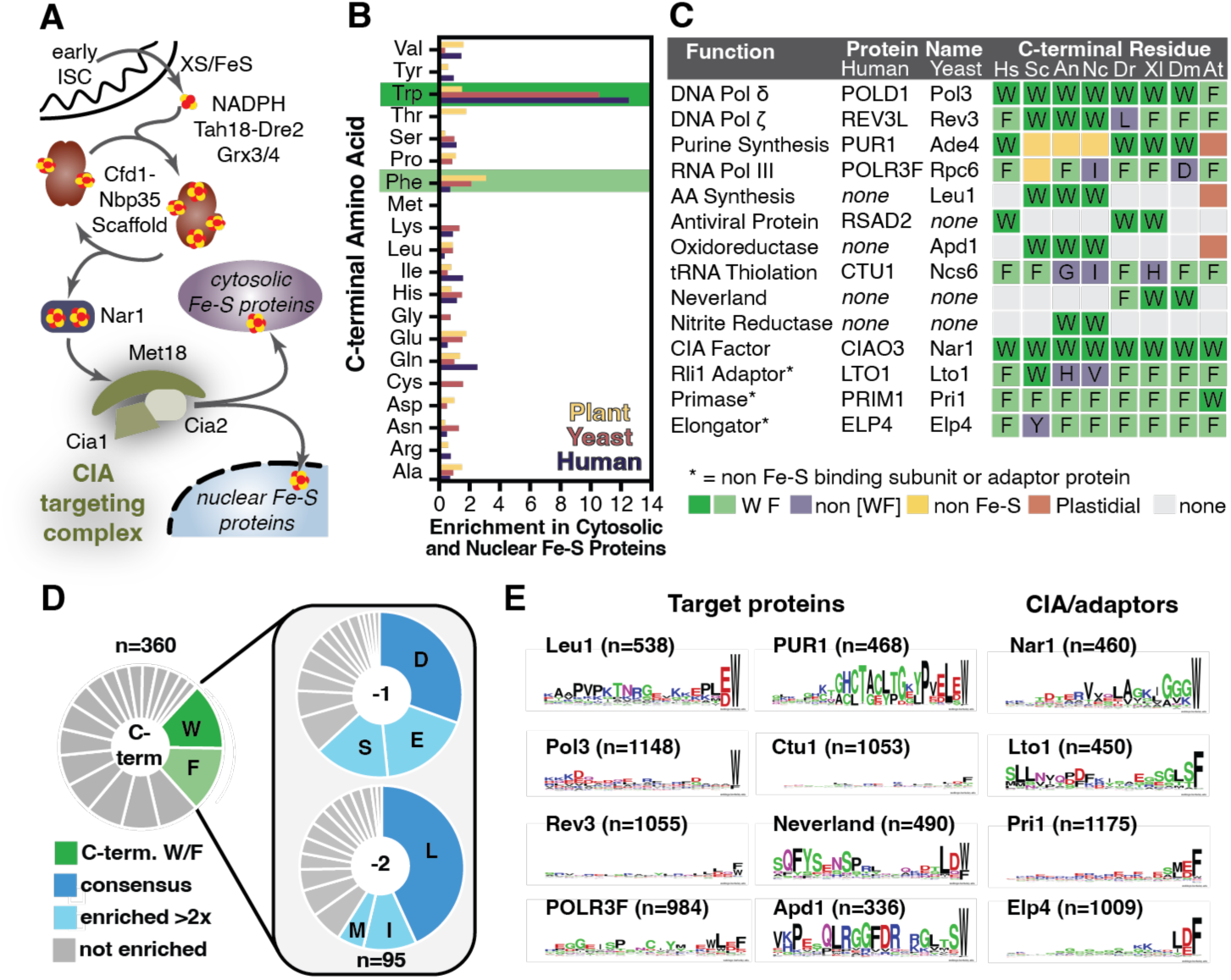
A conserved [LIM]-[DES]-[WF] tripeptide is found at the C-terminus of CIA clients. (**A**) Overview of the CIA pathway (yeast nomenclature). In the final step, the CIA targeting complex (CTC) identifies the CIA clients through direct, or adaptor mediated, protein-protein interactions and delivers the Fe-S cluster. (**B**) The enrichment of C-terminal amino acids (frequency in dataset/frequency in C-terminal proteome) in cytosolic and nuclear Fe-S proteomes of plants (*Arabidopsis thaliana*, yellow), yeast (*Saccharomyces cerevisiae*, pink), and in humans (*Homo sapiens*, purple). (**C**) The C-terminal residue of Fe-S proteins, CIA factors, and adaptors terminating in W/F in *H. sapiens* (Hs), *S. cerevisiae* (Sc), *Aspergillus nidulans* (An), *Neurospora crassa* (Nc), *Danio rerio* (Dr), *Xenopus laevis* (Xl), *Drosophilia melonogaster* (Dm), and *A. thaliana* (At). (**D**) Pie charts showing frequency of amino acids found at the C-terminus (n=360 sequences), and the −1 (penultimate) and −2 positions (n=95 sequences, those from the C-terminal dataset terminating in W/F). Amino acids indicated are enriched >2-fold (light green or blue) or >3-fold (dark green or blue) relative to their frequency in the human proteome. (**E**) WebLogos depicting conservation of C-termini for [4Fe-4S] proteins (Leu1, Pol3, Rev3, POLR3F, PUR1, Ctu1), [2Fe-2S] proteins (Neverland and Apd1), and the CIA factors/adaptors (Nar1, Lto1, Pri1, Elp4).

One major question addressed herein is how the CTC identifies its >30 clients, which include Fe-S proteins essential for DNA replication, transcription, translation, and other cellular processes (*19*). Proteomics and gene depletion studies have suggested that some clients only require a single CTC component for their maturation (*e.g*.Viperin (RSAD1) requires only Cia1), whereas other targets depend on two or more CTC subunits (*20, 21*). Additionally, the CIA client Rli1 does not directly bind the CTC but depends on the Yae1-Lto1 adaptor for CIA machinery recruitment (*22*). The diversity of the CIA proteome, combined with the incomplete catalog of CIA clients requiring adaptors, have made it challenging to unravel the cryptic codes driving substrate recognition by the CTC.

Despite these challenges, some recent studies indicated that some targets might exploit signals at their C-termini to recruit the CIA machinery. The clients, Viperin and Apd1, and the Lto1 adaptor terminate in a tryptophan residue that is critical for *in vivo* cofactor maturation (*22–24*). These studies suggest the tantalizing possibility that CIA clients use conserved recognition motifs to recruit the CTC, reminiscent of the recent proposal that an “LYR” tripeptide motif guides Fe-S cluster delivery from ISC (*8*). However, experimental support for this proposal is lacking, particularly regarding the number of eukaryotic Fe-S proteins bearing such signals, whether additional other residues in the primary or tertiary structure contribute to recognition, and whether these determinants are sufficient for CIA machinery recruitment. Here we show that >30% of CIA clients or their adaptors have a [LIM]-[DES]-[WF]-COO^-^ tripeptide motif that recruits the CIA machinery and demonstrate using integrative *in vitro* and *in vivo* approaches that this Targeting Complex Recognition (TCR) signal is both necessary and sufficient for binding the CTC and delivering Fe-S clusters from the CIA machinery.

## A C-terminal aromatic residue is enriched and conserved in CIA clients

By cataloging the C-terminal residue of every Fe-S protein in Fungi (*Saccharomyces cerevisiae*) and Metazoa (*Homo sapiens*) (*25*), we found that a C-terminal tryptophan residue was uniquely enriched in cytosolic and nuclear Fe-S proteins. Six (24%) of the yeast and five (13%) of the human CIA clients, factors, and adaptors terminated in a tryptophan (Fig. 1B, Data S1). For comparison, only 1.8% and 1.2% of all yeast and human proteins, respectively, end with a tryptophan. For plant (*Arabidopsis thaliana*) CIA clients (*26*), the most enriched C-terminal residue was phenylalanine, occurring in seven (17%) of 42 non-glutaredoxin cytosolic and nuclear Fe-S proteins (Fig. 1B, Data S1). Remarkably, <u>none</u> of the 164 yeast, human, or plant mitochondrial or plastidial Fe-S proteins terminated with a tryptophan or phenylalanine (fig. S1, Data S1). Similarly, just two (0.8%) of the 248 Fe-S proteins from Bacteria (*E. coli*) and Archaea (*Methanocaldococcus jannaschii)* terminate with tryptophan and ten (4%) with phenylalanine (fig. S1D, Data S1) (*27–29*). Therefore, a C-terminal tryptophan or phenylalanine is a hallmark of CIA clients – the first clear example of a putative CIA-targeting sequence.

Next, 360 CIA clients from 10 model organisms were analyzed challenge the universality of this hallmark and uncover any additionally conserved features (Fig. 1C, Data S2). Of the 95 sequences terminating in W or F (26% of 360 sequences), all had conserved amino acids in the penultimate (−1) and antepenultimate (−2) (Fig. 1D, Data S2). A negatively charged residue (48%) or a serine (15%) were preferred in the penultimate position, whereas a hydrophobic residue occurs at the antepenultimate in 59% of the sequences. Sequence alignments and analysis revealed conservation is limited to the −1 and −2 positions (Fig. 1E, Data S2). We named this tripeptide ([LIM]-[DES]-[WF]) motif the Targeting Complex Recognition (TCR) signal, due to its role in recruitment of the CTC (*vide infra*).

When CIA clients terminating in the TCR signal were compared to their homologs that are matured by other Fe-S biogenesis machineries, we only found the tripeptide motif to be exclusively conserved in CIA clients. For example, fungal isopropylmalate isomerases (Leu1) are cytosolic [4Fe-4S] proteins with a TCR signal (Fig. 1E), but bacterial, archaeal, mitochondrial and plastidial orthologs are missing the tripeptide (fig. S2, A and B). Analogous correspondences occur for other classes of Fe-S proteins, including: the CIA factor Nar1; the [4Fe-4S] protein Viperin, Ctu1, and nitrite reductase; and the [2Fe-2S] proteins Apd1, Neverland, and choline monooxygenase (fig. S2). The length of the TCR-tail varies between ∽3 and 45 amino acid residues (fig. S2, boxed), suggesting the tail length tunes positioning of different clients. Available proteomic mass spectrometry data and structures of Fe-S proteins isolated from the native host demonstrates that the TCR motif remains attached after maturation (table S2), in contrast to mitochondrial import.

Additionally, some Fe-S clusters play a structural or regulatory role and therefore their cluster binding residues can be lost with evolutionary drift. Remarkably, the TCR signal and Fe-S ligands show strong correlation. For example, many glutamine phosphoribosylpyrophosphate amidotransferases, such as PUR1 in humans, are [4Fe-4S] proteins and they terminate with a TCR signal (Fig. 1E). In contrast, TCR signals are absent in the plant enzymes, which are plastidial, and thus are not matured by CIA, and in their fungal homologs, which lack Fe-S clusters (fig. S3A). RNA polymerases displayed similar co-conservation of TCR and Fe-S ligands (fig. S3B). These correlations strongly suggest that the TCR signal is functionally related to CIA.

Next, we truncated the TCR motif at the C-terminus of Pol3, the catalytic subunit of DNA polymerase δ (*30*). Pol3 is essential for yeast viability upon DNA damage with methylmethane sulfonate (MMS) (Fig. 2A). Chromosomal *POL3* was placed under control of a galactose-regulatable promoter (Gal-*POL3*). This strain’s phenotypic growth defect in the presence of MMS and glucose was alleviated by episomal Pol3 expression at endogenous levels, but not by modified genes encoding truncated Pol3 variants (Fig. 2A). In similar experiments, C-terminally truncated variants of Apd1, Nar1, and Leu1 were unable to restore normal growth when expressed at low levels in Gal-regulatable (GAL-*NAR1*) or deletion (Δ*apd1*, Δ*leu1*) strains (Fig. 2, B to D; fig. S4 and S5). In all cases, removal or substitution of just the C-terminal tryptophan was sufficient to elicit the growth defect (Fig. 2). Thus, when a the TCR motif is present, it is universally required for the Fe-S protein function.

**Fig. 2.**
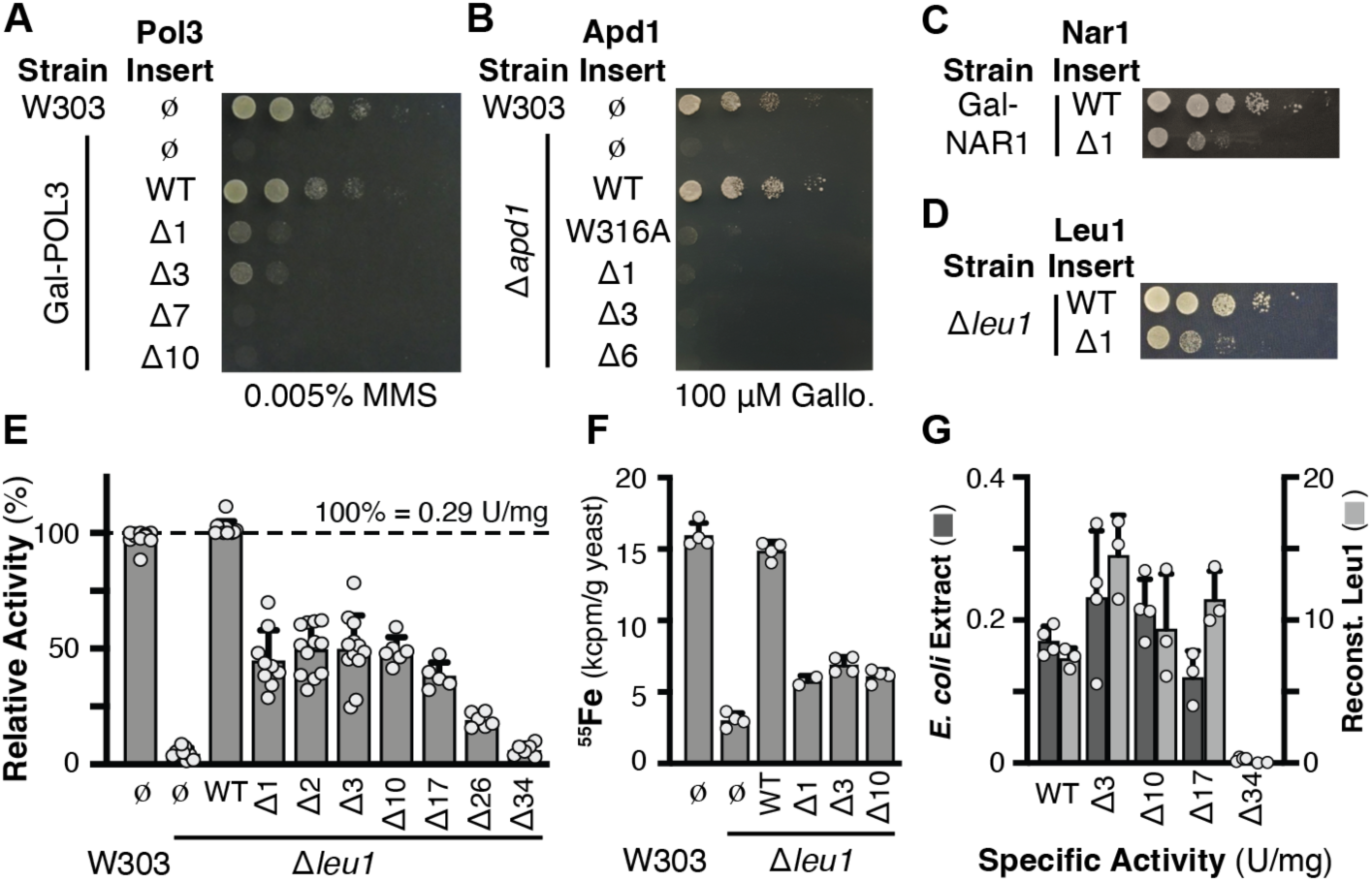
Deletion of the C-terminal tryptophan of CIA clients results in defective Fe-S cluster delivery. (**A**-**C**) The indicated yeast strains, wild-type W303, *apd1* deletion, or Gal-regulable *POL3* or *NAR1*, were transformed with a centromeric plasmid for expression of Pol3 (A), Apd1 (B), or Nar1 (C) from their natural promoters. The Δ indicates the number of C-terminal amino acids that were removed, ø indicates the empty vector control, and WT indicates the full length wild-type insert. Yeast cells were grown overnight in glucose (Glc) containing medium, then spotted after serial dilution onto Glc medium with the indicated concentrations of methylmethane sulfonate (MMS) or gallobenzophenone (Gallo). (**D**) Plasmid-borne Leu1 was expressed from the weak (*RET2*) promoter in a Δ*leu1* strain with a functional *LEU2* allele. Yeast were grown as in (A-C), but in the absence of leucine. (**E** and **F**) Determination of Leu1 enzymatic activity in yeast crude extracts and ^55^Fe incorporation to quantify Fe-S cluster insertion into Leu1. W303 or Δ*leu1* strains harboring the indicated Leu1 variant (expressed from its natural promoter) were grown as in (A-C). For *de novo* ^55^Fe incorporation cells were grown in Fe-free medium overnight, followed by incubation with ^55^Fe and immunoprecipitation of Leu1. (**G**) Fe-S cluster insertion by the *E. coli* ISC machinery (cell extract, dark gray) or by chemical reconstitution of Leu1’s Fe-S cluster following purification (light gray) are unaffected by C-terminal truncation up to 17 amino acids.

Focusing on Leu1, the benchmark enzyme for quantitative measurements of CIA function (*31*), centromeric plasmids encoding Leu1 truncations were expressed in a Δ*leu1* strain. Deletion of W779 from the TCR (Leu1-Δ1), and a further truncation up to 17 amino acids, led to 60 % loss of Leu1 activity without decreasing the Leu1 expression level. (Fig. 2E; fig. S6). Analysis of *de novo* ^55^Fe incorporation into Leu1 was similarly diminished by TCR truncation (Fig. 2F). We speculate that the remaining activity of Leu1-Δ1 results from a second CTC binding determinant (*32*). At low expression levels, both the TCR and this second determinant are required for efficient maturation, but increased expression can drive CTC-Leu1-Δ1 or -Nar1-Δ1 complex formation (Fig. 2, C and D; fig. S4). Additional controls demonstrated the decreased Leu1-Δ1 activity was independent of the transcriptional promoter and terminator sequences (fig. S7). To rule out the possibility that the Leu1 activity loss results from defective protein folding, the Leu1 truncations were expressed in *E. coli*, which does not encode the CIA machinery. Leu1 activity in *E. coli* crude extracts or following its purification and Fe-S cluster reconstitution was undiminished upon removal of ≤17 amino acids (Fig. 2G). These data establish that the Leu1 activity loss upon TCR signal truncation results from a specific defect in cluster delivery from the CIA system.

## The TCR motif mediates interaction with the CIA Targeting Complex

The CIA targeting complex (CTC) identifies apo-clients and thus it is the likely TCR signal receptor. To test this hypothesis, we used an affinity copurification assay to analyze the TCR-CTC interaction, employing the [2Fe-2S] Apd1 and [4Fe-4S] Leu1 client proteins. Both proteins copurified with the CTC, but variants lacking the indole moiety had a defect in CTC binding (Fig. 3, A to C; fig. S8). Thus, the TCR tail is critical for CTC interaction *in vitro*, explaining why TCR truncation leads to a defective Fe-S protein maturation *in vivo*.

**Fig. 3.**
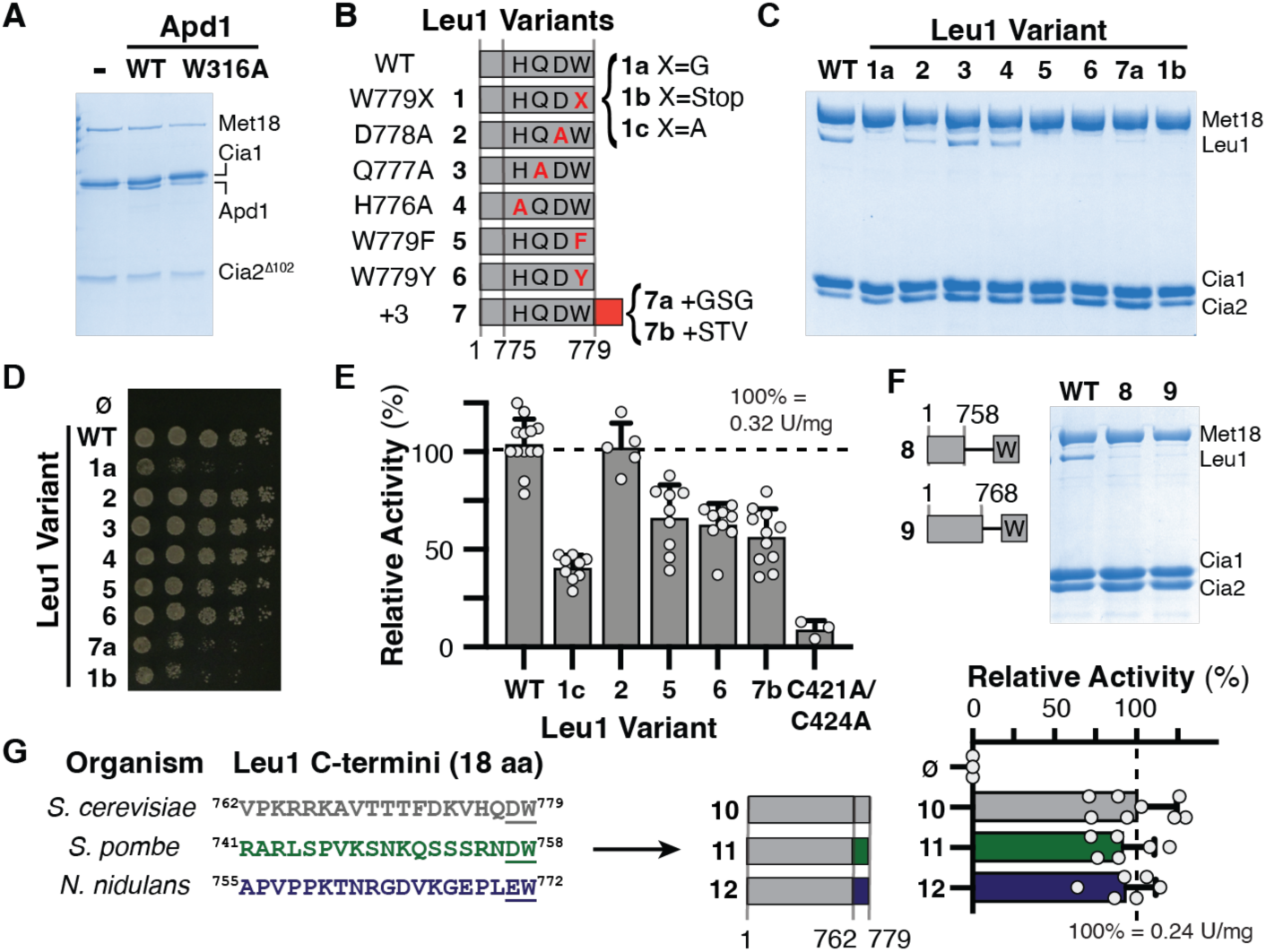
CTC binding *in vitro* and Fe-S cluster maturation *in vivo* depend on the C-terminal tryptophan and its carboxy terminus. (**A**) Apd1 (wild-type; WT; W316A variant, or none; -), was mixed with yeast CTC subunits (Met18, Strep-tagged Cia1, and truncated Cia2^Δ102^). Apd1 associated with the CTC was evaluated via SDS-PAGE analysis after Streptactin affinity purification. (**B)** Variants (red) of wild type (WT) Leu1 tested in panels (C-E). Non-bold numbers refer to the positions in the amino acid sequence. (**C**) The Leu1 variants (bolded numbers) were mixed with the yeast CTC (Met18, Cia1, and Strep-tagged Cia2). Leu1 associated with the CTC was evaluated by SDS-PAGE after Streptactin affinity purification. (**D**) Leucine-independent growth of yeast (as in Fig. 2D) depends on the C-terminal W of Leu1. (**E**) Determination of Leu1 variant enzymatic activity in yeast cell extract as in Fig. 2E. (**F**) Affinity copurification of C-terminal Leu1 truncations as in (B). (**G**) The last 18 amino acids of *S. cerevisiae* Leu1 (gray, **10**) were replaced with Leu1 tails from *S. pombe* (green, **11**) or *A. nidulans* (purple, **12**). The specific activities of the resulting tail transplant variants were analyzed as in Fig. 2E.

To confirm the functional relevance of the TCR to client protein maturation *in vivo*, the Leu1 W779 variants were expressed at low levels in a Δ*leu1* strain to evaluate growth in a leucine deficient medium and at higher levels to quantify Leu1 activity in crude extracts (Fig. 3, D and E; fig. S9). The W779A complemented strain grew at a 2-fold slower rate than strains expressing the WT protein and exhibited a 2-fold reduction in Leu1 specific activity (Fig. 3, D and E; fig. S9). Western blotting confirmed the decreased activity was not due to differences in protein levels (fig. S10). Leu1 with W779F/Y substitutions did not pull down with the CTC in the stringent *in vitro* assay (Fig. 3C), but the *in vivo* functionality of these TCR variants was less severely affected (Fig. 3, D and E; fig. S9). Altogether, these results demonstrate that a C-terminal tryptophan is the preferred residue for yeast TCR signals, but TCRs that include a phenylalanine residue can also be recognized.

Next, we extended the C-terminus of Leu1 by three amino acids to determine whether an internal TCR signal retained functionality. This construct was clearly unable to bind the CTC or recruit the CIA machinery *in vivo* (Fig. 3). Since the D778A, Q777A, and H776A substitutions neither impacted yeast growth nor CTC binding (Fig. 3), our data identify the aromatic side chain <u>and</u> its C-terminal position as the most critical determinants for CTC recruitment.

To examine whether TCR tail length impacts its functionality, we replaced the last 10 or 20 amino acids of Leu1 with a single tryptophan residue. As neither variant copurified with the CTC (Fig. 3F), we concluded that either additional amino acids of the tail are critical for recognition by the CTC or the TCR signal has a certain spatial requirement to allow for correct positioning in the CTC complex. To distinguish between these possibilities, we replaced the last 18 amino acids of the *S. cerevisiae* Leu1 with those from *Schizosaccharomyces pombe* or *Aspergillus nidulans* proteins. Importantly, these three sequences do not share any conserved residues besides the terminal [DE]-W motif (Fig. 3G). Because these tail transplant variants were efficiently matured by the CTC (Fig. 3G), we concluded that in addition to the primary sequence determinants, the TCR tail length impacts client positioning for optimal interaction and Fe-S delivery from the CTC.

## The TCR signal is sufficient for recruitment of CTC to non-native targets

To eliminate any additional sequence determinants within CIA clients that might contribute to CTC recruitment, we fused various TCR tails to SUMO, a small well folded protein with an accessible C-terminus. These SUMO peptide carriers (SPC) for Leu1_759-779_, Nar1_482-491_, Pol3_1088-1097_ and Rev3_1495-1504_ each bound to the CTC in a tryptophan dependent manner (Fig. 4A). Moreover, the Leu1 tail could be further truncated from 20 to just 3 amino acids (QDW) without disrupting CTC interaction (Fig. 4A).

**Fig. 4.**
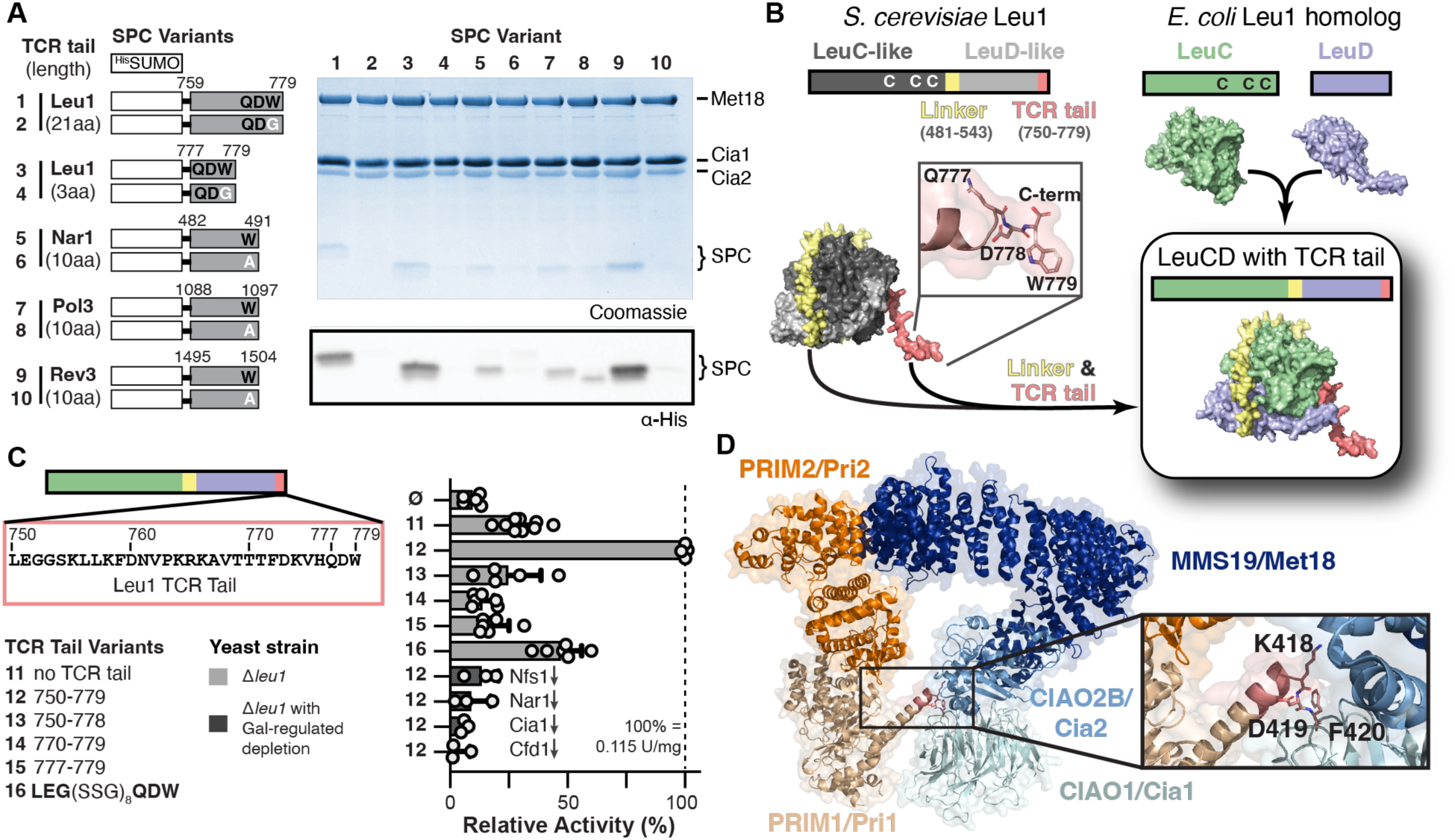
An engineered TCR signal is sufficient to recruit non-native proteins to the CTC and for Fe-S cluster delivery. (**A**) Cartoon of SUMO peptide carrier (SPC) fusions used in copurification, as in Fig. 3C. Streptactin elution fractions were analyzed by SDS-PAGE and immunoblotting against the SPC N-terminal His-tag. (**B**) A non-native Fe-S protein, *E. coli* LeuCD, was engineered for cluster delivery from the CIA machinery. The *E. coli* subunits were fused with a linker (yellow) and the TCR-tail (salmon) of yeast Leu1 was attached. Protein structures are AlphaFold models. (**C**) *In vivo* cytosolic [4Fe-4S] cluster insertion into engineered *E. coli* LeuCD depends on the TCR signal with an appropriate tail length. Specific activities in cell extracts were determined as in Fig. 2E. Each LeuCD variant was expressed in a Δ*leu1* yeast strain (light gray), or a Δ*leu1* strain with galactose-regulatable genes (dark gray). Growth on Glc medium for 40 h led to depletion (↓) of the indicated Fe-S cluster biosynthesis protein. (**D**) Modelling of the PRIM1-PRIM2 (AlphaFold, orange and sand) and CTC (6TC0, blue hues) into the CryoEM density of the Primase-CTC complex (*32*). This model shows that the TCR signal (KDF, salmon) interacts with the proposed client binding site on blade 3 of CIAO1.

If the TCR motif is sufficient for association with the CTC, then it should be possible to direct Fe-S cluster delivery to a non-native, heterologous Fe-S protein in yeast. To test this prediction, we focused on the *E. coli* LeuC-LeuD heterodimer, the homolog of yeast Leu1 (Fig. 4B). Because the [4Fe-4S] cluster of LeuC-LeuD is biosynthesized by the bacterial ISC system, it does not possess the TCR-extension (fig. S2, A and B). The *E. coli leuC* and *leuD* genes were fused with linker derived from yeast Leu1 (yellow, Fig. 4B). When the last 30 amino acids of yeast Leu1 were fused to the C-terminus of this LeuCD construct, its enzymatic activity increased >3-fold (Fig. 4C). To demonstrate this increased activity was due to CIA machinery recruitment, several control experiments were performed. First, we determined that the increased activity of LeuCD-TCR fusion depends on the C-terminal tryptophan (Fig. 4C). Second, fractionation of yeast demonstrated LeuCD activity is cytosolic, not mitochondrial where ISC is functional (fig. S11). Third, GAL-regulatable expression of Fe-S assembly factors in τι*leu1* strains confirmed the increased activity depends on the CIA machinery and the early ISC machinery providing XS/[Fe-S]_int_ (Fig. 4C).

To identify the minimal length required to position LeuCD for cluster delivery, we shortened the TCR tail to 3 or 10 amino acids, but these variants were unable to guide Fe-S maturation (Fig. 4C). However, when Leu1 tail residues 753-776 were replaced with (SSG)_8_ spacer, 47±9% activity was recovered (Fig. 4C), corroborating our observation that the TCR tail length is critical for proper positioning (Fig. 3, F and G). Thus, the TCR signal can be exploited to guide non-native Fe-S proteins to the CIA machinery for metallocofactor delivery.

## Discussion

N- or C-terminal primary sequence determinants are important signals for subcellular trafficking and post-translational modifications (*33*). Herein, we establish that cytosolic and nuclear Fe-S protein maturation is another key process controlled through short linear peptide motifs presented on the solvent-exposed C-terminus. The [LIM]-[DES]-[WF] tripeptide is ubiquitous in the eukaryotic domain of life, and consistent with our proposal it serves as a universal signal directing cytosolic and nuclear Fe-S protein maturation. Since CIA is proposed to biosynthesize [4Fe-4S] proteins (*1, 34*), we thus were initially surprised to identify TCR motifs in four bis-histidinyl coordinated [2Fe-2S] proteins, including Apd1 which requires its C-terminal tryptophan for CTC interaction and *in vivo* function (Fig. 2B, Fig. 3A). Consequently, our data imply that [2Fe-2S] proteins are also matured by the CTC.

Surprisingly, we uncovered “cryptic” TCR motifs in ELP4 and PRIM1 (Fig. 1E, fig. S12), which are non-metal binding subunits of the elongator and primase complexes. We speculate ELP4 and PRIM1 serve as adaptors for ELP3 and PRIM2, the [4Fe-4S] subunits, similar to Lto1-Yae1 for Rli1 maturation (*22*). First, PRIM1’s TCR signal is well conserved in Eukaryotes, but absent in archaeal homologs (fig. S12), following the pattern observed for other TCRs (fig. S2). Second, PRIM1 was found to be required for formation of the primase-CTC complex (*32*), but the molecular basis for this requirement was puzzling. By docking of PRIM1 into the available cryoEM density of the CTC-PRIM1-PRIM2 complex (*32*), we discovered that PRIM1’s TCR signal is properly positioned to interact with the CIAO1 and CIAO2 subunits of the CTC (Fig. 4D). This previously unnoted finding not only corroborates our proposal that PRIM1 can function as an adaptor for PRIM2, but it indicates that the TCR motif of clients and adaptors anchors in this region of the CTC (*32, 35*). Altogether, our data explain how ∼30% of clients are identified by the CIA machinery: 19% (6 of the 32 human CIA clients) directly interact with the CTC via their TCR, and an additional 9% (ELP3, PRIM2 and Rli1) depend on an adaptor.

One important question remains unresolved: how do the remaining cytosolic and nuclear Fe-S proteins, those without TCR signals or adaptors, recruit the CTC? Our findings combined with previous data indicates that at least two TCR determinants are present in every CIA client and the successful approach described herein provides a framework for discovery of these additional motifs. Finally, it is notable that just one determinant is sufficient to recruit the CIA machinery, as the [LIM]-[DES]-[WF] motif can be exploited to engineer cluster delivery to a non-native Fe-S enzyme (Fig. 4). As deficiencies in cluster maturation frustrate efforts to engineer novel biosynthetic pathways requiring Fe-S enzymes, our study provides a roadmap for overcoming these bottlenecks and demonstrates how an improved understanding of the fundamental biochemistry can impact metalloenzyme bioengineering.

## Acknowledgments

We thank Sheena Vasquez for assistance in generating the model shown in Figure 4D that was based on a cryo-EM map deposited by Kassube and Thomä. We are grateful to Roland Lill’s lab, particularly Martin Stümpfig, for ^55^Fe incorporation analysis. We also thank Gary Sawers, Sean Elliott, Sheena Vasquez, Catherine Drennan, and Dan Kahne for helpful discussions.

## Author contributions

Conceptualization: DLP, AJP

Methodology and Investigation: MDM, CG, AB, YL, CB, SB, LL, AH, KW, KN, DJN

Visualization: MDM, DLP, AJP

Funding acquisition: DLP, AJP

Project administration: DLP, AJP

Supervision: DJN, DLP, AJP

Writing – original draft: MDM, DJN, DLP, AJP

Writing – review & editing: MDM, AB, AH, DJN, DLP, AJP

## Data and materials availability

All data are available in the main text or the supplementary materials and materials used in the analysis are available upon request any researcher for purposes of reproducing or extending the analysis.

## Supplementary Materials for

### Materials and Methods

#### SUMO peptide carrier (SPC) plasmid construction and purification

The SPC variants were constructed using the pTB146 plasmid (gift from Thomas Bernhardt, Harvard University). DNA encoding the C-terminal tail of Leu1 (FDNVPKRKAVTTTFDKVHQDW; QDW), Nar1 (KDLVSVGSTW), Pol3 (QEKVEQLSKW) and Rev3 (EEALISLNDW) were inserted at the 3’-end of the His-SUMO coding region in pTB146 via Q5 Site-Directed Mutagenesis Kit (NEB) or by gene synthesis (for SPC-Nar1 and SPC-Pol3, GENEWIZ). Plasmids were transformed into an *E. coli* BL21(DE3). Cells were grown at 37°C until an OD_600_ 0.7, isopropyl β-D-1-thiogalactopyranoside (IPTG, 0.5 mM) was added, and cells were harvested 4 hours later. For purification, cells (∼10 g wet cell paste) were resuspended in 100 mL of Buffer A [50 mM Tris-HCl (pH 8.0), 100 mM NaCl, 5% glycerol, 5 mM ϕ3−mercaptoethanol (ϕ3ME)] supplemented with 1 mM PMSF (GoldBio), 4 kU DNase nuclease (Fisher Scientific), 5 mM imidazole, and 1 mg/mL lysozyme (GoldBio). Cells were disrupted by sonication and extract clarified by centrifugation. Extract was applied to a 4 mL Ni-NTA resin. The column was washed with >15 column volumes (CV) of Buffer A containing 20 mM imidazole and eluted with Buffer A with 350 mM imidazole. Protein containing fractions were combined, and exchanged into Buffer A via dialysis or gel filtration, concentrated to 10-20 mg/mL via ultrafiltration (Amicon, 10 kDa cutoff filter, Millipore), and stored at −80°C.

#### Apd1 cloning and purification

The plasmid with Apd1 cloned into pDONR201 was obtained from the plasmid repository at Arizona State University (DNASU) and inserted via Gateway cloning into the pDEST17 vector, a commercially available gateway destination plasmid for N-terminally His-tagged expression in *E. coli* to create the His-tagged Apd1 (^His^Apd1) expression plasmid. The W316A substitution was introduced via Q5 Site-Directed Mutagenesis Kit (NEB) (table S3). All sequences were confirmed by DNA sequencing. For expression, plasmids were transformed into an *E. coli* BL21(DE3) with the ISC operon (a gift from Markus Ribbe,(*36*)) and cells were grown and induced as previously described (*24*). For purification, typically 5 g of cell paste was resuspended in 50 mL of Buffer B (50 mM Tris-HCl (pH 9.0), 300 mM NaCl, 5% glycerol, 5 mM ϕ3ME) supplemented with 1 mM PMSF (GoldBio), one protease inhibitor tablet (Thermo Scientific), 4 kU DNase nuclease (Fisher Scientific), and 5 mM imidazole. Cells were lysed by addition of CelLytic Express (2.5 g, Sigma Aldrich) and extract was clarified by centrifugation. After batch absorption for 1h, the Ni-IMAC resin (GoldBio) was collected, washed with 20 CV of Buffer B with 20 mM imidazole, and eluted in Buffer B with 250 mM imidazole. Protein containing fractions were combined, exchanged into storage buffer [50 mM Tris-HCl (pH 9.0), 100 mM NaCl, 5% glycerol, 5 mM ϕ3ME], concentrated to 4-7 mg/mL via ultrafiltration (Amicon, 10 kDa cutoff filter, Millipore), and stored at −80°C.

#### Affinity Co-purification Analysis

Leu1 variants in Fig. 3, C and F were created in previously described Leu1 plasmid encoding a TEV-cleavable N-terminal His-tag (^HisTEV^Leu1) (*21*). ^HisTEV^Leu1 variants were created by Q5 mutagenesis (NEB, table S3) and confirmed by DNA sequencing. IMAC purification of ^HisTEV^Leu1 was carried out as described (*21*). In the co-purification assay, Met18 with a N-terminal His-SUMO tag (^SUMO^Met18); singly tagged Cia1 with an N-terminal His-tag (^His^Cia1) or double tagged Cia1 (^DT^Cia1) with an N-terminal His-TEV-Strep tag; doubly tagged Cia2 (^DT^Cia2) with a thrombin-cleavable N-terminal Strep-tag and a HRV3C-cleavable C-terminal His-tag; and truncated Cia2 (Cia2^Δ102^) in which the first 102 aa in Cia2’s N-terminus and a C-terminal His-tag, were purified as previously described (*21, 37*). When required, the His-tag was removed from ^DT^Cia1 via treatment with TEV protease and isolation of resulting singly Strep-tagged Cia1 (^Strep^Cia1).

Affinity copurification assays were carried out as previously described (*21*). Briefly, a Strep-tagged bait protein, ^DT^Cia2 (for Leu1 and SPC-fusions) or ^Strep^Cia1 (for Apd1), was mixed with an equimolar amount of each CTC subunits (^SUMO^Met18, ^His^Cia1 or Cia2^Δ102^, as appropriate) and a TCR-tail prey protein (Leu1, Apd1 or SPC-fusions). For Apd1 experiments, the truncated Cia2 (Cia2^Δ102^) was used due to the close molecular weights of the full length Apd1, ^Strep^Cia1, and Cia2. For SPC fusions, roughly 3-fold molar excess over bait protein concentrations were used. In all experiments, a “no-bait” negative control was completed in parallel to detect any nonspecific binding to the resin. The samples were incubated for 1 h 4°C and then chromatographed through streptactin resin. Input and Elution fractions were analyzed SDS-PAGE (12% for experiments with Leu1, 15% for SPC-fusions, 20% for Apd1). For Western blot analysis, proteins were transferred using a Trans-Blot Turbo Mini PVDF Transfer pack and Transfer System with the mixed molecular weight protocol in the manufacturer’s instructions (Bio-Rad). The membrane was blocked with 5% casein in TBST and probed with Anti-His antibody (Cell Signaling Technology). Bands were detected via chemiluminescence, according to manufacturer’s instructions.

#### Yeast Strains and Cell Growth

The *Saccharomyces cerevisiae* strain W303-1A (MATa, *ura3-1*, *ade2-1*, *trp1-1*, *his3-11,15*, *leu2-3,112*) was used as the wild-type strain. Yeast strains are summarized in table S4. To convert the W303-1A strain into a *LEU2*^+^/Δ*leu1* strain, first homologous recombination with a *leu2* PCR fragment followed by selection on SC glucose medium lacking leucine was carried out. The *leu2* PCR product was obtained by fusion PCR of *leu2* fragments A, B and C (see table S5). Yeast strains, Δ*leu1*, *LEU2*^+^/Δ*leu1*, Gal-NFS1/Δ*leu1*, Gal-*NAR1*/Δ*leu1*, Gal-CIA1/Δ*leu1*, and Gal-CFD1/Δ*leu1*, were constructed by homologous recombination in which the coding region of the *leu1* gene was replaced by a natNT2 cassette amplified from pYM17, omitting the 6xHA tag (*38*). Gal-*POL3* was constructed by homologous recombination in which the nucleotide-upstream promoter region was replaced by the natNT2 cassette from pYM-N27 (*38*), including a GAL promoter sequence. The correctness of the insertion was assured by PCR analysis of genomic DNA. Cells were grown on rich (YP) or minimal (SC) media containing 2 % (w/v) galactose or glucose as carbon source with the appropriate markers.

#### Yeast expression plasmid construction

For *POL3* cloning, the *E. coli* strain NEB5 F’I^q^ was used. All other genes were cloned using the *E. coli* strains NEB5 or NEB10. *E. coli* genomic DNA was used as the template to amplify *leuC* and *leuD*. For the other genes (*POL3, NAR1, LEU1, APD1*), *S. cerevisiae* genomic DNA was the template for PCR. Information, including primers (table S6) and plasmids (table S7), are provided and when not specified otherwise, standard cloning procedures were used. Variants were generated by site-directed mutagenesis with primer design according to Zheng *et al.* (*39*). For C-terminal truncations, the appropriate codon was replaced by a stop codon. Successful construction of all plasmids was verified by Sanger sequencing.

The *POL3* coding region and the 500 bp at the 5’ end of the start codon was inserted into the SacI and EcoRI sites of pRS416-MET25_prom_-MCS-CYC1_term_ (maintaining the *CYC1* terminator) (*40*). During cloning and mutagenesis of *POL3*, growth for plasmid minipreps required incubation of agar plates at room temperature for 2 days until colonies appeared and cultivation in liquid media at 30 °C.

For *APD1*, its coding region, the 526 bp natural promoter, and 327 bp terminator regions were cloned into the SacI and KpnI sites of pRS416-MET25_prom_-MCS-CYC1_term_.

The *NAR1* coding region, along with its natural promoter (591 bp) and terminator (558 bp) were cloned into SacI and KpnI sites of pRS416-MET25_prom_-MCS-CYC1_term_. For the Nar1-encoding plasmid under control of the *MET25* promoter, the *NAR1* coding region and its terminator (558 bp) were cloned into SpeI and KpnI sites of pRS416-MET25 (*40*). For the Nar1 plasmid with *NBP35* promoter, the promoter region of the *NBP35* gene (508 bp) was inserted between the SacI and SpeI sites of pRS416-MET25_prom_-Nar1-NAR1_term_.

The *LEU1* coding region, along with its natural promoter (552 bp) and terminator (512 bp) regions, were cloned as two PCR fragments employing the natural SalI site (883–888) in the *LEU1* coding region. First, the SalI-NgoMIV fragment was cloned into pRS416-MET25_prom_-MCS-CYC1_term_, followed by the SacI-SalI fragment. Variants with other promoter regions were generated by introducing a SpeI site just before the ATG start codon of *LEU1*, followed by cloning PCR fragments of the promoter regions of the *MET25* (412 bp), *TDH3* (649 bp), *RET2* (500 bp) and *RPL18B* (500 bp) genes between SacI and SpeI sites. For pRS416-TDH3_prom_-Leu1-TDH3_term_, an XhoI site was introduced immediately following the *LEU1* stop codon and a BamHI site was inserted after the *LEU1* terminator. Then, a PCR fragment of 449 bp corresponding to the *TDH3* terminator was cloned into the XhoI and BamHI sites.

For generation of pRS416-TDH3_prom_-Leu1-TDH3_term_ with the C-terminal 19 amino acids of the yeast Leu1 replaced by the C-terminal 19 amino acids of the homologs of *S. pombe* or *A. nidulans,* a BglII was introduced by mutagenesis at nucleotides 1847-1852 of the *leu1* gene, creating a silent mutation. Then, synthetic genes corresponding to the BglII sequence plus nucleotides 1853 to 2280 of yeast *LEU1*, and yeast codon optimized nucleotides encoding the 19 C-terminal amino acids of the homologs plus a stop codon and the XhoI sequence were cloned into BglII/XhoI sites of pRS416-TDH3_prom_-Leu1-TDH3_term_.

For construction of the yeast expression vector for LeuCD (pRS426-FBA1_prom_-LeuC-linker-LeuD-Leu1-CT-FBA1_term_), *E. coli leuC* and *leuD* genes were each amplified and separately cloned into the SpeI/XhoI sites of the MCS of pRS424 and pRS426, respectively. To create the backbone of the shuttle vector, the *TDH3* promoter of vector pRS426-TDH3_prom_-MCS-CYC_term_ (*40*) was replaced by 1000 bp of the *S. cerevisiae FBA1* promoter, using SacI and SpeI sites. Then, the *CYC1* terminator was exchanged for the *FBA1* terminator (1000 bp), using XhoI and KpnI sites. To create pRS426-FBA_prom_-LeuD-Leu1-TCR, the DNA fragment encoding the yeast Leu1 C-terminus (last 30 amino acids, Leu1-TCR) was cloned into EcoRI/BamHI of the multiple cloning site of pRS426-FBA1_prom_-MCS-FBA1_term_ followed by cloning of *leuD* from *E. coli* (using pRS426 with *E. coli leuD* as template) into pRS426-FBA1_prom_-Leu1-TCR-FBA1_term_. To introduce *leuC* and the linker region of Leu1 (amino acids 481-543) concomitantly, PCR overlap extension was used (*41*). In the first PCR reaction (I), *leuC* (in pRS424) was amplified such that the forward primers introduced a SpeI restriction site and the reverse primer introduced 15 nucleotides corresponding to the 5’ end of the *LEU1* linker. In the second PCR reaction (II), the *LEU1* linker was amplified (from pRS416-*TDH3*_prom_-Leu1-*TDH3*_term_). The forward primer introduced 20 nucleotides corresponding to the 3’end of *leuC* and the reverse primer appended an SpeI restriction site. An equimolar amount of PCR-product, I and II, were mixed and hybridized by using a PCR protocol without dNTPs. In a final PCR reaction, the primers flanking the SpeI sites were used to amplify the product, which was subsequently cloned into the SpeI restriction site of pRS426-FBA_prom_-LeuD-Leu1-TCR, generating pRS426-FBA_prom_-LeuC-LeuD-Leu1-TCR-FBA_term_. To test the effect of the Leu1-TCR, a stop codon was introduced after the *leuD* coding region by site-directed mutagenesis. To create the QDW tail, primers were used to insert the tripeptide coding region plus a stop codon immediately following *leuD*.

To shorten the TCR tail from 29 to 10 amino acids (770-779 amino acids of Leu1), Q5 mutagenesis was used to delete the region corresponding to amino acids 750-769 of Leu1-TCR and install an Xho1 site. The resulting PCR product was digested with XhoI cut and ligated. For the construct with the SSG linker sequence substituting the Leu1-TCR, a synthetic DNA sequence encoding for the eight SSG repeats and the codons for QDW was chemically synthesized and cloned into the XhoI/EcoRI sites of 426-FBA1_prom_-LeuC-linker-LeuD-Leu1-TCR-FBA1_term_.

#### Protein expression and purification of Leu1 variants for activity measurements

Yeast Leu1 was cloned into pET28a (Novagen), fused to a N-terminal hexa-histidine tag for heterologous expression and purification. After transformation into BL21 (DE3) cells (NEB), an overnight pre-culture was diluted (2% inoculum) into 2 liters of LB medium containing ampicillin and 3% (v/v) ethanol p. a and allowed to grown at 37 °C and 200 rpm/min shaking. When the culture reached an OD_600_ of 0.5, the induction was started by addition of IPTG (0.5 mM), followed by incubation at overnight at 30 °C. Cells were opened with French Press and purified using Ni-IDA (Cube) according to the manufacturer’s instructions. The protein eluates were desalted using a Sephadex G-25 column (GE Healthcare) equilibrated with 25 mM Tris-HCl, pH 8.0, 300 mM NaCl. The purified proteins were shock-frozen and stored at −80 °C until use.

Reconstitution of recombinant purified apo-Leu1 was performed in an anaerobic chamber (Coy Laboratory). Approximately 50 µM of Leu1 was reduced with DTT (15 mM, end concentration) for 45 min. After addition of 5 molar-equivalents of ammonium iron (III) citrate and development of a red color the same equivalents of lithium sulfide were slowly pipetted into the reaction tube. The sample was incubated for 2-4 h and the Leu1 activity was tested during this time without desalting the sample.

^55^Fe incorporation into Leu1

Leu1 or mutants thereof cloned into pRS416 plasmids under the control of the endogenous *LEU1* promoter were transformed into yeast cells, as indicated in figure captions. *In vivo* radiolabeling with ^55^FeCl_3_ and determination of ^55^Fe incorporation into Leu1 beads was carried out as described (*31*). Briefly, freshly transformed cells were grown on iron-poor SC medium for 40 h supplemented with the appropriated carbon source. After incubation with ^55^FeCl_3_ and 1 mM ascorbate for 2 h in SC iron-poor medium, cell extracts were prepared by using glass beads. The Leu1 protein was immunoprecipitated by using polyclonal antibodies against Leu1 bound to Protein A-Sepharose beads. The amount of protein-associated radioactivity was assessed via liquid scintillation counting.

#### Growth complementation

Growth complementation assays used the W303, Gal-*POL3*, Δ*apd1*, Gal-*NAR1* and *LEU2*^+^/Δ*leu1* strains transformed with appropriate plasmids (see figure legends). Transformants were grown in liquid SC medium supplemented with galactose or glucose (2 %) at 30 °C for 40 h (Δ*apd1*, *LEU2*^+^/Δ*leu1* and Gal-*NAR1*, including control W303), or for 16 h (series Gal-*POL3*, including control W303). At this point, the cultures were diluted in SC medium to an OD_600_ of 0.5 and 5 µL aliquots were submitted to a 10-fold serial dilution and spotted into SC agar plates supplemented with glucose or galactose carbon source. SC agar plates with methyl methane sulfonate or gallobenzophenone at the indicated concentration were freshly prepared. The plates were incubated for 48 h at 30 °C and photographed. At least three experiments using independently transformed yeast cells were performed.

#### Enzyme activity determination

Protein determination for activity in crude extracts was performed using the Microbiuret method with desoxycholate/trichloroacetic acid coprecipitation (*42, 43*). Isopropylmalate isomerase and succinate dehydrogenase activity were determined in freshly prepared yeast or *E. coli* cell extracts. Yeast cell lysates were prepared using the glass bead method and clarified by centrifugation, whereas *E. coli* cell lysates were prepared using a French Press. For Leu1 activity, the reaction was conducted in a buffer containing 20 mM Tris-HCl, pH 7.4 and 50 mM NaCl. The formation of isopropylmaleate after addition of 0.2 mM 3-isopropylmalate was followed by the increase of absorption at 235 nm. To correct for variations in growth conditions and deviations from the ambient temperature (22 °C), a wild type Leu1 or LeuC-linker-LeuD-TCR (variant 12, Fig. 4C) control was measured in parallel and used to normalize the series measured on different days.

For the coupled assay of Leu1 activity, *E. coli* isopropylmalate dehydrogenase *(leuB*) was cloned into the BamHI and HindIII sites of MCS1 of pET-Duet1 and purified by Ni-NTA using standard procedures. Assays contained 20 mM Tris-HCl (pH 8), 50 mM KCl, 10 mM MgCl_2_, 1 mM DTT, 0.2 mM isopropylmaleate, 0.4 mM NAD^+^, 2 mM pyrazol and 0.4 U LeuB. The absorbance increase from NADH formation at 340 nm was followed.

The succinate dehydrogenase activity in isolated mitochondria was determined by measuring the reduction of cytochrome *c* at 550 nm in a reaction containing 8.9 mM succinate and 1 mg/mL bovine heart cytochrome *c* in a buffer containing 50 mM Tris, pH 8.0 and 50 mM NaCl. The background activity was assayed by measuring the increase of absorbance in the same reaction containing 14.5 mM malonate and 1 mM potassium cyanide.

#### Bioinformatic analysis

Fe-S proteins and adaptor amino acid sequences were collected by NCBI BlastP searches using yeast und human sequences as query. Amino acid sequence alignment was carried out with ClustalO on the EMBL-EBI server (*44*). The same server was used for the download of reference proteomes for the analysis of C-termini (Data S1). For analysis of the C-termini of *Homo sapiens* (*25*), *Saccharomyces cerevisiae* (*19*), *E. coli* (*28*), *Methanocaldococcus jannaschii* (*29*), and *Arabidopsis thaliana* (*26*) the published Fe-S protein inventories were manually updated. Datasets for the generation WebLogos (*45*) were downloaded by selection of the appropriate taxonomic class of organisms in the OrthoDB catalogue of orthologs (*46*). Sequence fragments lacking N-terminal or C-terminal regions and excessively long sequences due to erroneous translation and/or wrong assignment of introns were manually removed. Then the unaligned C-terminal 20 amino acids were submitted to the Weblogo server for data in Fig. 1E, and fig. S3. The Apd1 collection was from the phylogenetic tree as described in Stegmaier et al.(*24*).

**Fig. S1.**
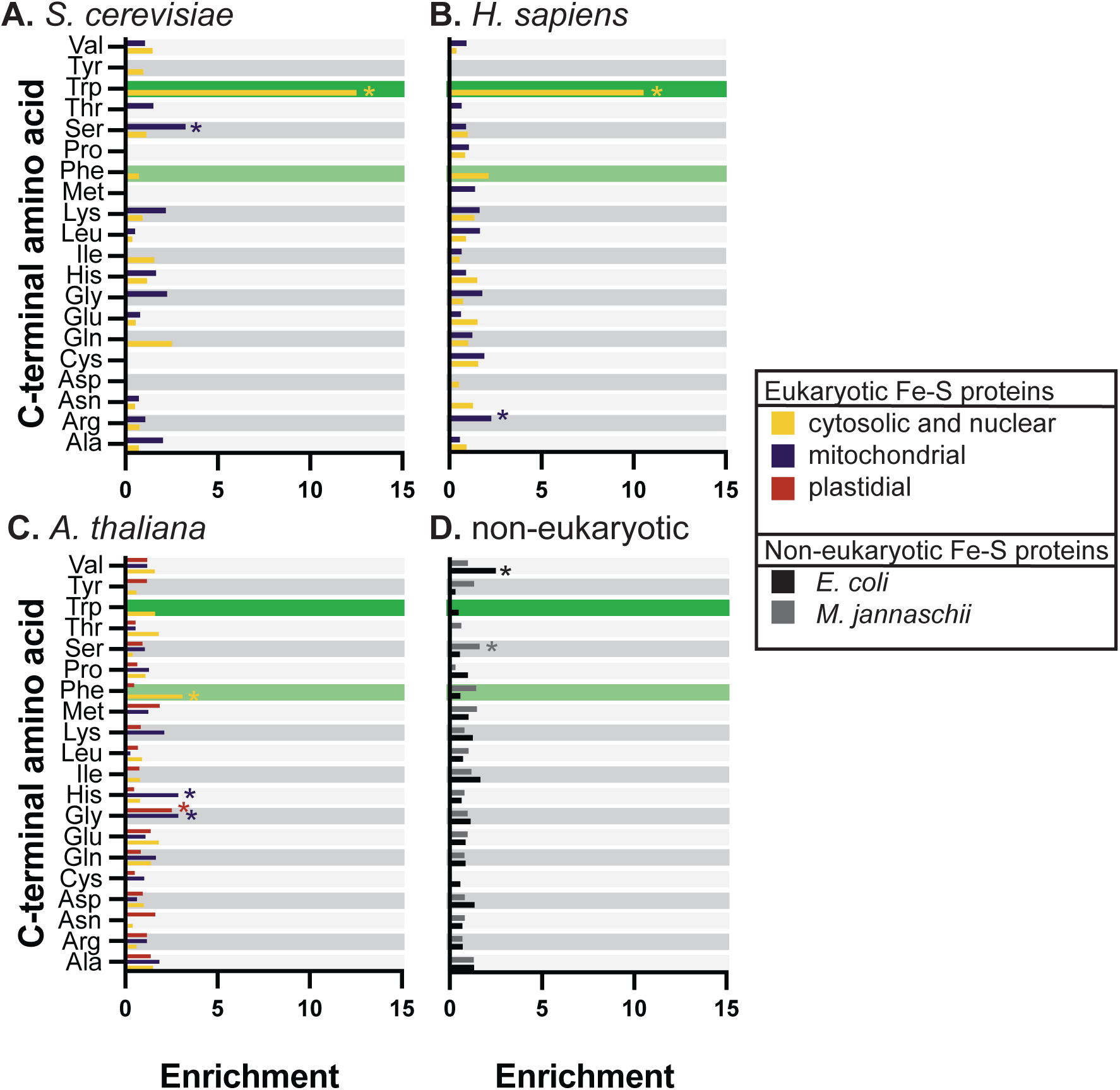
Bar graphs comparing enrichment in C-terminal amino acids in the Fe-S proteomes of yeast (**A**), humans (**B**), plants (**C**), and non-eukaryotic organisms (**D**). For Panels (A-C), the CIA proteome is in yellow, the mitochondrial Fe-S proteome is purple, and plastidial Fe-S proteome is red. In Panel D, the *Methanocaldococcus jannaschii* Fe-S proteome is in light gray and *Escherichia coli* Fe-S proteome is black. In each dataset, the C-terminal amino acid with the highest enrichment is marked with an asterisk. A terminating W (dark green) or F (light green) is exclusively enriched in the CIA proteome.

**Fig. S2.**
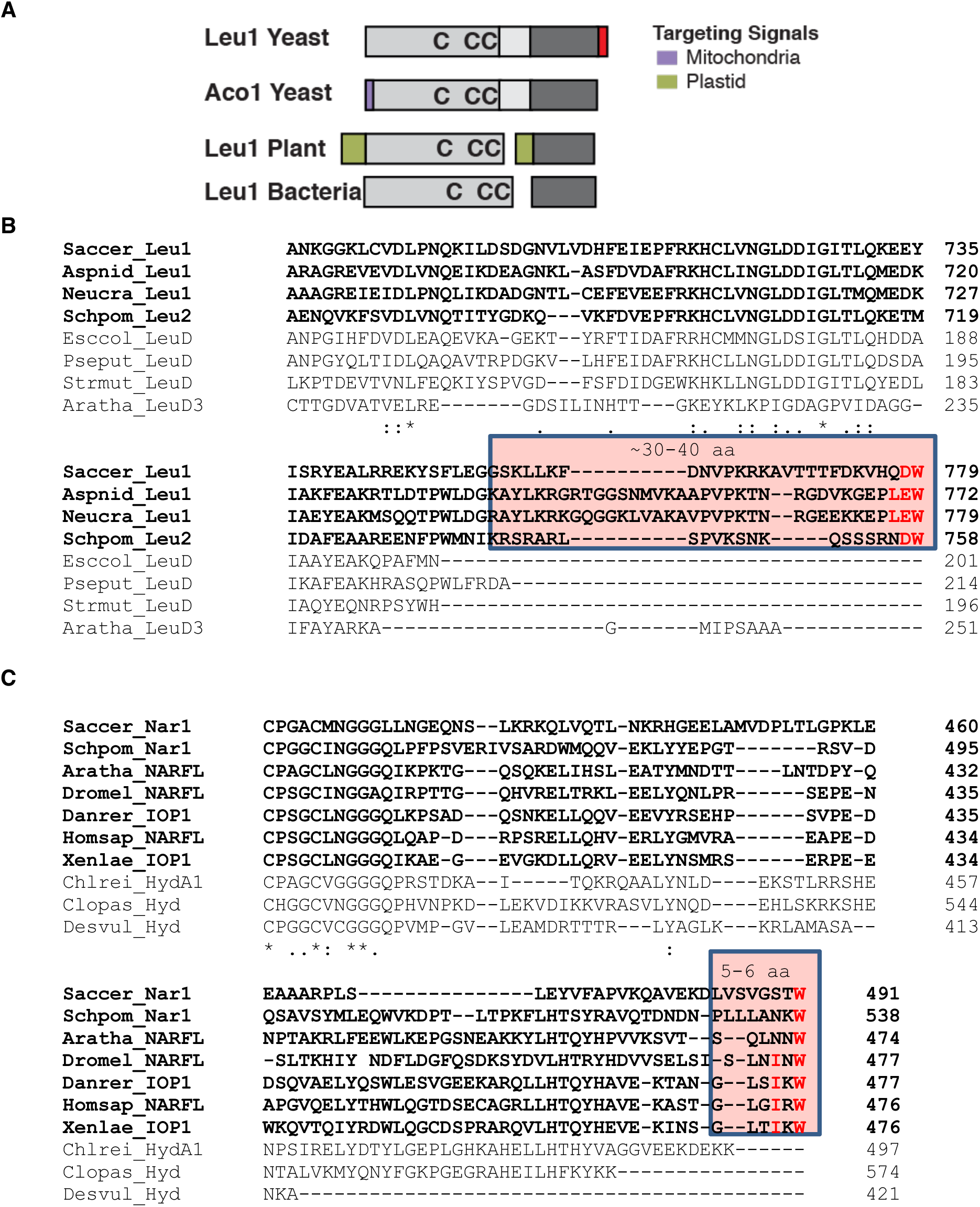
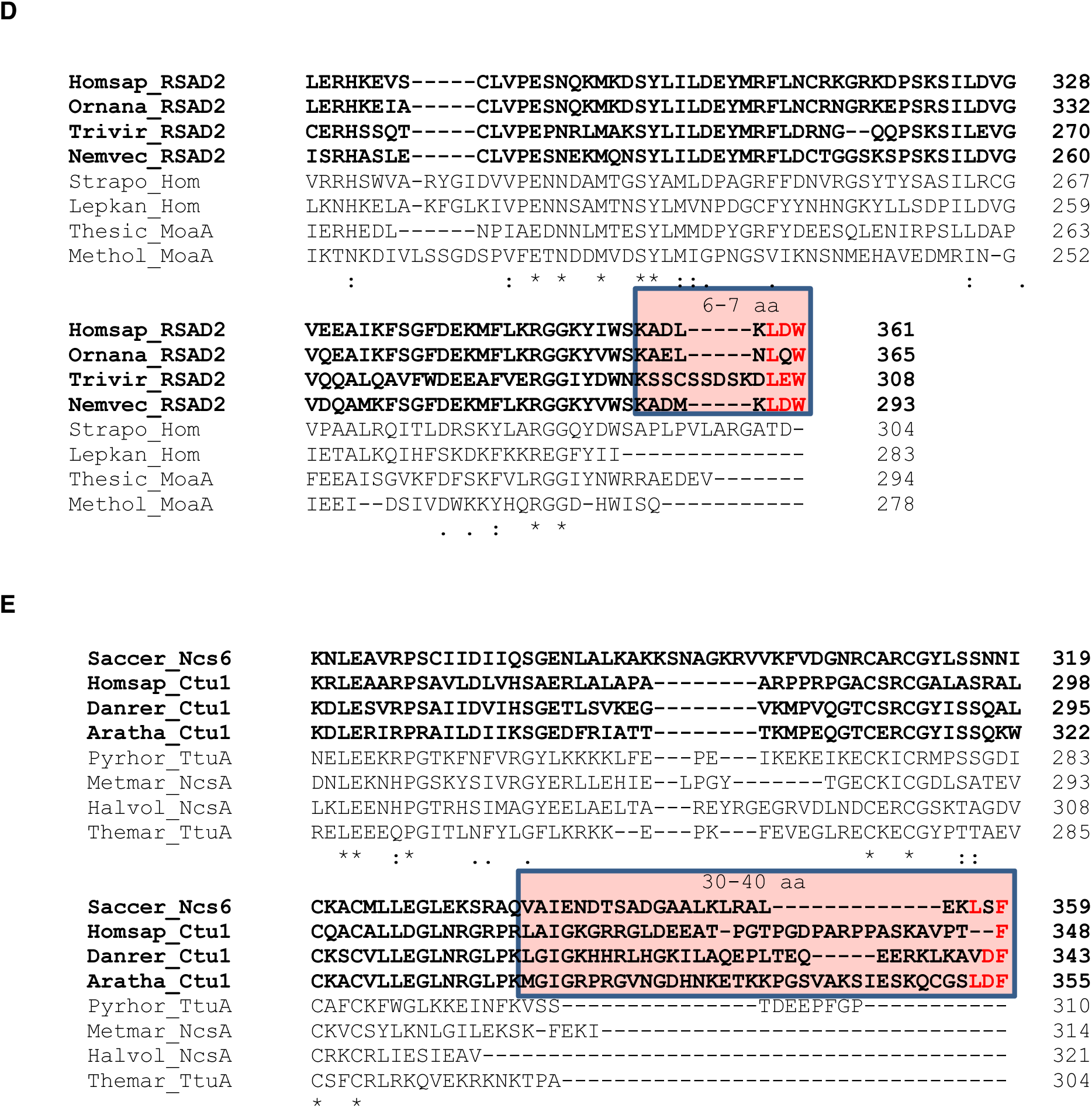
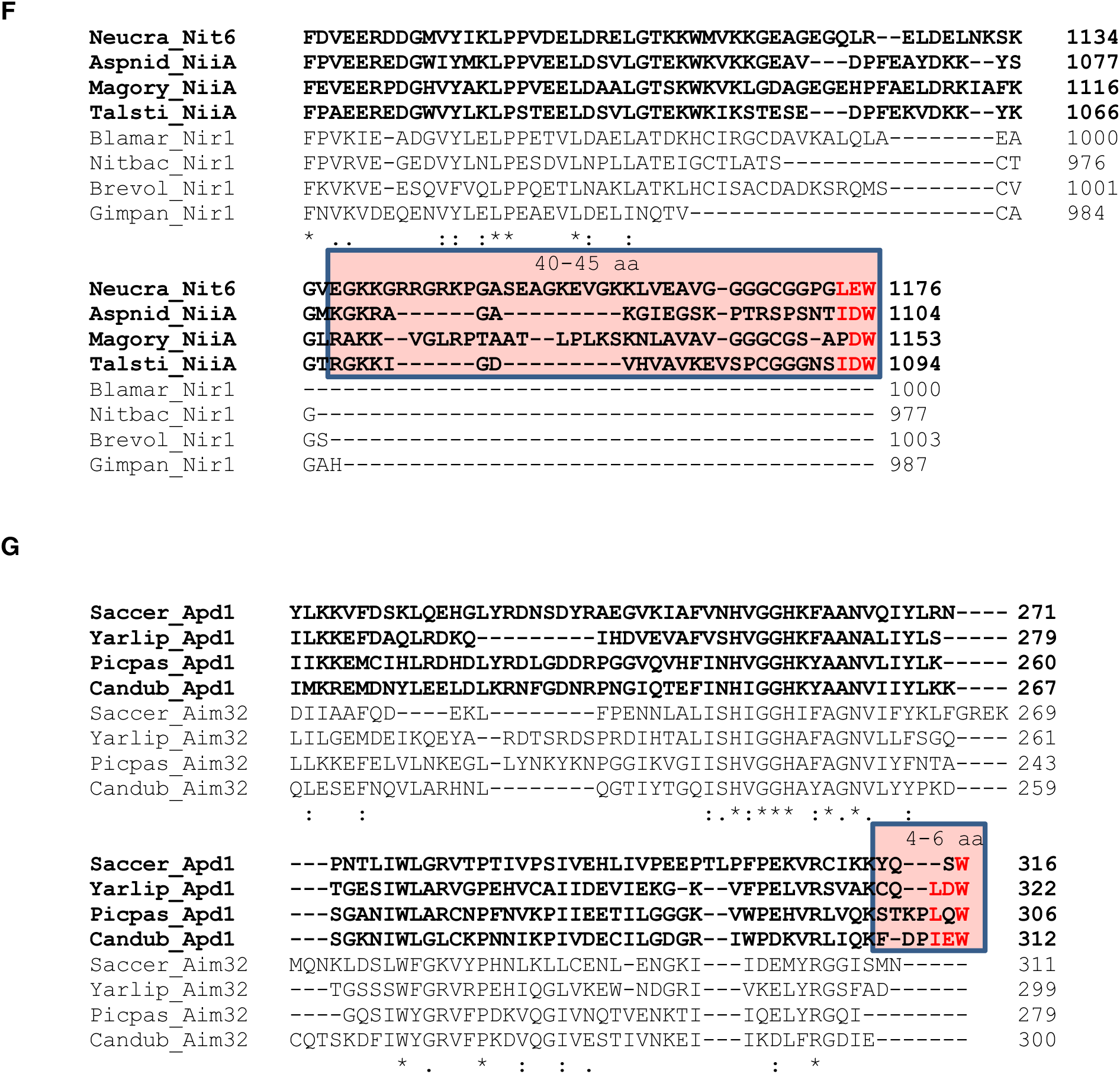
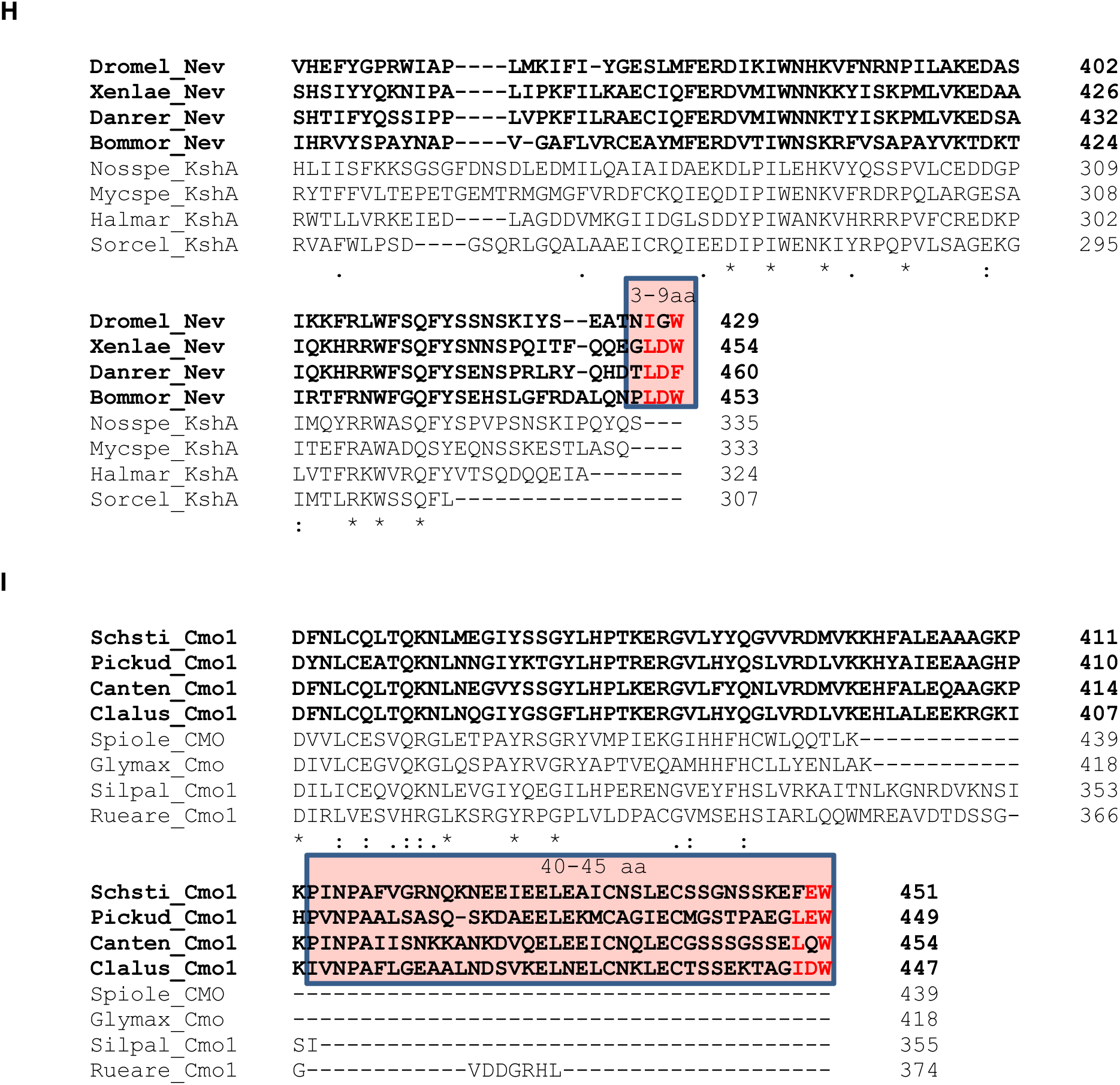
A terminating in W/F is conserved in CIA substrates (bold) at the end of a C-terminal extension (boxed with length of extension noted), but not in eukaryotic homologs with organellar targeting sequences, nor in Bacterial or Archaeal homologs. (**A**) Cartoon of isopropylmalate isomerase and aconitase in Fungi, and isopropylmalate isomerase subunits in Plants and Bacteria. (**B**). Amino acid sequence alignment of C-termini of fungal Leu1 sequences (Saccer, *Saccharomyces cerevisiae*; Aspnid, *Aspergillus nidulans*; Neucra, *Neurospora crassa*; Schpom, *Schizosacharomyces pombe*, historically called Leu2) compared with bacterial LeuD (Esccol, *Escherichia coli*; Pseput, *Pseudomonas putida*; Strmut, *Streptococcus mutans*, PDB 2HCU) and plastidial LeuD3 (Aratha, *Arabidopsis thaliana*). (**C**) Amino acid sequence alignment of C-termini of eukaryotic Nar1/IOP1/NARFL sequences (Saccer, *Saccharomyces cerevisiae*; Schpom, *Schizosaccharomyces pombe*; Aratha, *Arabidopsis thaliana*; Dromel, *Drosophila melanogaster*; Danrer, *Danio rerio*; Homsap, *Homo sapiens*; Xenlae, *Xenopus laevis*) and Fe-Fe hydrogenases (Chlrei, *Chlamydomonas reinhardtii*; Clopas, *Clostridium pasteurianum*; Desvul, *Desulfovibrio vulgaris*). (**D**) Amino acid sequence alignment of C-termini of eukaryotic Viperins (Homsap, *Homo sapiens*; Ornana, *Ornithorhynchus anatinus*; Trivir, *Trichoderma virens*, PDB 7N7I; Nemvec, *Nematostella vectensis*, PDB 7N7H) compared with bacterial (Strapo, *Streptomyces apocyni*; Lepkan, *Leptospira kanakyensis*) and archaeal homologs (Thesic, *Thermococcus siculi*; Methol, *Methanomethylovorans hollandica*). (**E**) Amino acid sequence alignment of C-termini of eukaryotic Ncs6/Ctu1 proteins (Saccer, *Saccharomyces cerevisiae*; Homsap, *Homo sapiens*; Danrer, *Danio rerio*; Aratha, *Arabidopsis thaliana*) with archaeal homologs (Pyrhor, *Pyrococcus horikoshii*, PDB 5MKQ; Metmar, *Methanococcus maripaludis* S2, PDB 6SCY; Halvol, *Haloferax volcanii* DS2; Themar, *Thermotoga maritima* MSB8). (**F**) Amino acid sequence alignment of C-termini of fungal Nit-6/NiiA nitrite reductases (Neucra, *Neurospora crassa* OR74A; Aspnid, *Aspergillus nidulans* FGSC A4; Magory, *Magnaporthe oryzae* 70-15; Talsti, *Talaromyces stipitatus* ATCC 10500) and their archaeal Nir1 homologs (Blamar, *Blastopirellula marina*; Nitbac. *Nitrospirales* bacterium, HIB54509.1; Brevol, *Bremerella volcania*; Gimpan, *Gimesia panareensis*). (**G**) Amino acid sequence alignment of C-termini of fungal Apd1 (cytosolic) and Aim32 (mitochondrial) homologs (Saccer, *Saccharomyces cerevisiae*; Yarlip, *Yarrowia lipolytica*; Picpas, *Pichia pastoris*; Candub, *Candida dublinensis*). (**H**) Amino acid sequence alignment of C-termini of Neverland, eukaryotic Rieske-type hydroxylases (Dromel, *Drosophila melanogaster*; Xenlae, *Xenopus laevis*; Danrer, *Danio rerio*; Bommor, *Bombyx mori*) and bacterial ketosteroid hydroxylase homologs (Nosspe, *Nostoc sp*. NIES-4103; Mycspe, *Mycobacterium sp*. Aquia_213; Halmar, *Halioglobus maricola*; Sorcel, *Sorangium cellulosum*). (**I**) Amino acid sequence alignment of C-termini of Rieske-centre containing fungal choline monoxygenases (Schsti, *Scheffersomyces stipitis* CBS 6054; Pickud, *Pichia kudriavzevii*; Canten, *Candida tenuis* ATCC 10573; Clalus, *Clavispora lusitaniae* ATCC 42720) with plant plastidic (Spiole, *Spinacia oleracea*; Glymax, *Glycine max*) and bacterial homologs (Silpal, *Silvanigrella paludirubra*; Rueare, *Ruegeria arenilitoris*).

**Fig. S3.**
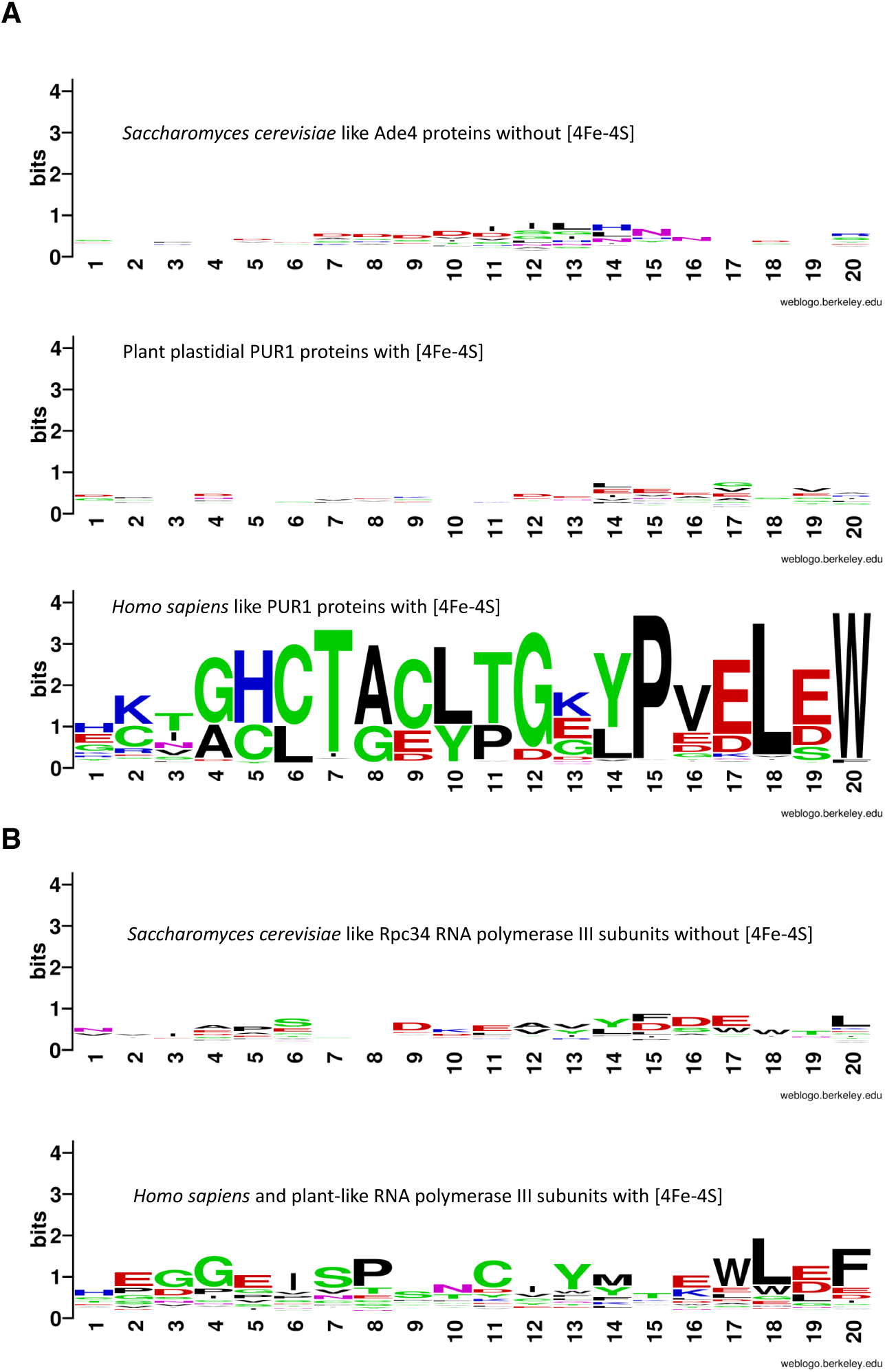
Conservation of a C-terminal W/F correlates with conservation of Fe-S cluster binding ligands. (**A**). Absence (top two Weblogos) and presence (bottom) of a C-terminal W or F residue correlates with the absence and presence, respectively, of Fe-S cluster binding signatures in eukaryotic glutamine phosphoribosylpyrophosphate amidotransferases (PUR1). Weblogo depictions for the C-terminal 20 amino acids of 570 yeast-like Ade4 proteins, which do not bind a [4Fe-4S] cluster, of 205 plant PUR1 proteins, which due to their plastidial localization lack a C-terminal W/F signature, and of 468 human-like PUR1 proteins, which based on conservation of 4 cysteine residues bind a [4Fe-4S] cluster. Sequences were collected via OrthoDB (*46*) and sorted according to their amino acid sequence identities with yeast, human and plant PUR1/Ade4 proteins and manually annotated for the Fe-S binding ligands. (**B**). Absence (top) and presence (bottom) of a C-terminal W or F residue correlates with the absence and presence, respectively, of Fe-S cluster binding signatures in eukaryotic RNA polymerases. WebLogo depictions for the C-terminal 20 amino acids of 131 yeast-like (Rpc34) RNA polymerase III subunits, which do not bind a [4Fe-4S] cluster and of 984 human-like (POLR3F) RNA polymerase III subunits, which based on conservation of 4 cysteine residues bind a [4Fe-4S] cluster. Sequences were collected via OrthoDB. The presence of a cysteine residue corresponding to Cys307 in the human sequence in a multiple amino acid sequence alignment was used to define sequences predicted to bind a [4Fe-4S] cluster.

**Fig. S4.**
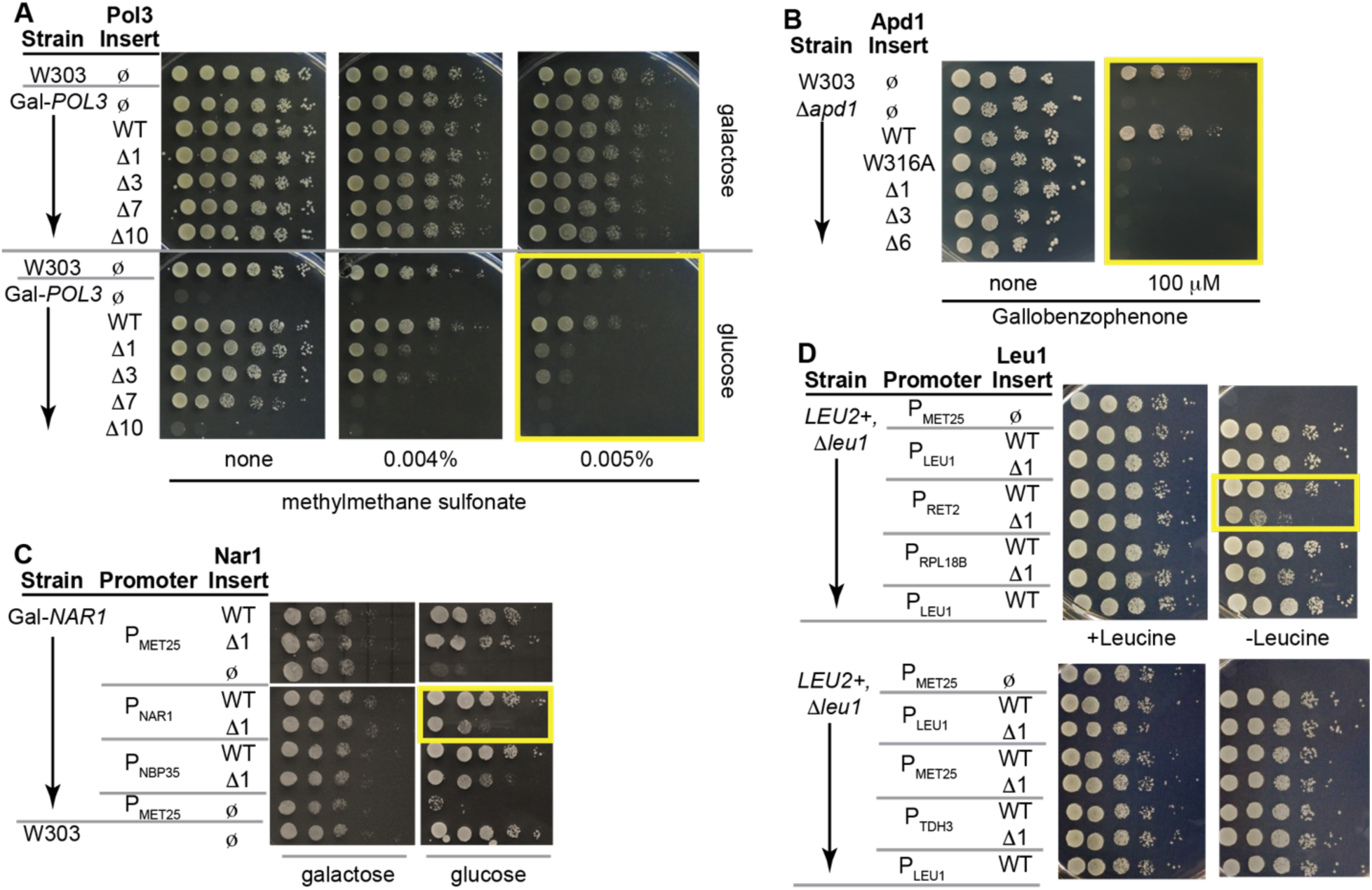
Non-cropped images of droptests shown in Figure 2A-D (yellow boxes illustrate data in main text) along with controls showing growth under permissive conditions including: in the presence of glucose (**A** and **C**); in the presence of leucine (**D**); or in the absence of gallobenzophenone (**B**). For Nar1 (C) the strong *MET25* or the weaker *NBP35* and *NAR1* promoters were used to drive expression of the WT and Δ1 inserts. For Leu1 (D) and Nar1 (C), the weak (*RET2, NBP35* and *RPL18B*), moderate (*LEU1*) and strong (*MET25* and *TDH3*) promoters were used. In both cases, the stronger promoters can mask the growth defect observed for the Δ1 variants in comparison to the weaker promoters.

**Fig. S5.**
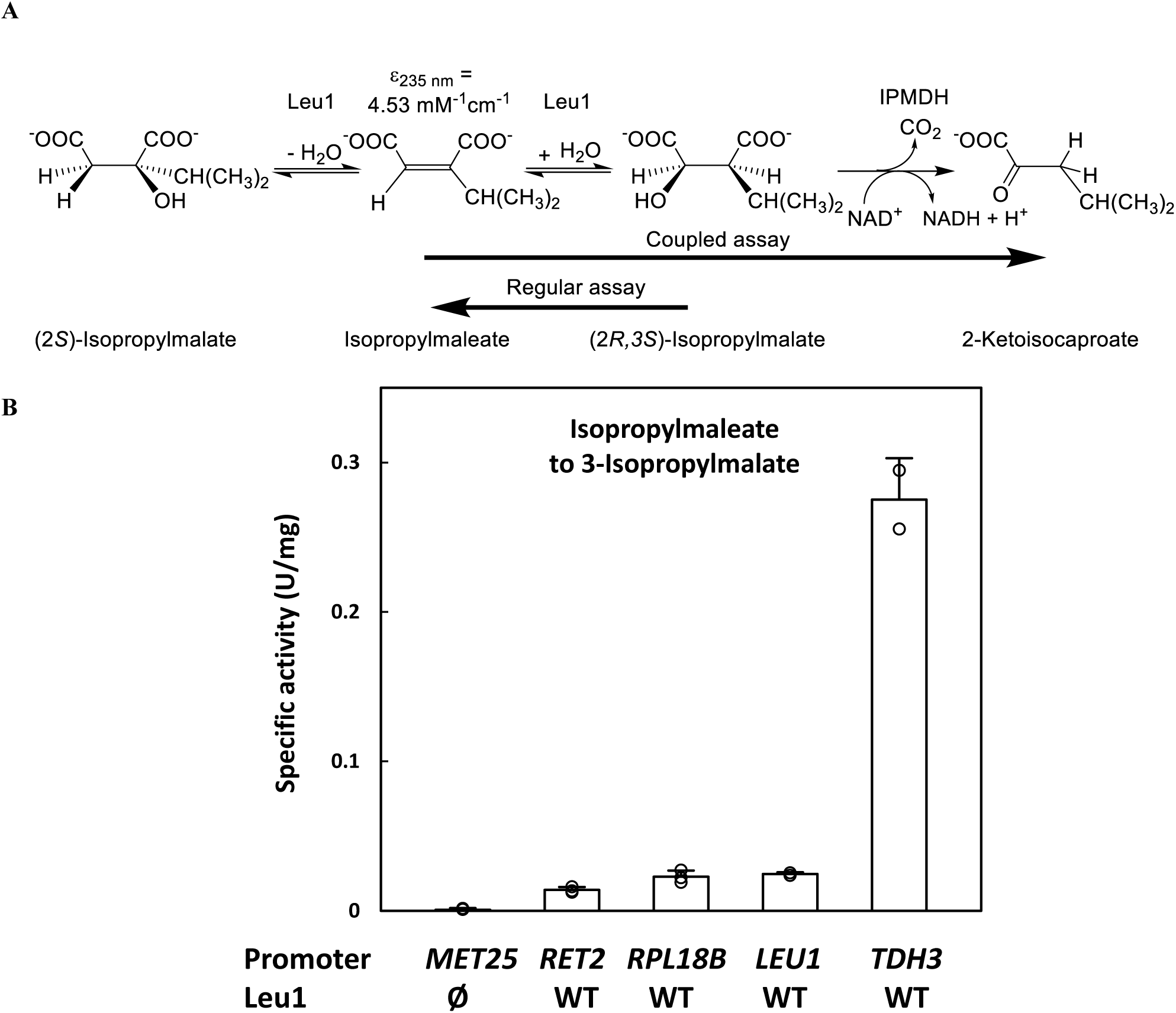
(**A**) The regular (*31*) and coupled isopropylmalate isomerase assay. (**B**) Bar graph with specific activities of isopropylmalate isomerase in cell extracts from Δ*leu1*/*LEU2^+^*cells transformed with pRS416 plasmids with *LEU1* under control of indicated promoters. Cells were grown in SC glucose medium including leucine. A coupled assay (3-isopropylmalate formation from isopropylmaleate as detected by *E. coli* isopropylmalate dehydrogenase (IPMDH) dependent NADH formation was used to quantify the isopropylmalate isomerase activity in cell extracts with *RET2*, *RPL18B*, *LEU1* and *TDH3* promoter driven *LEU1* constructs. Under these growth conditions the use of a *MET25* promoter for *LEU1* expression results in 70±6 % (n=3) of the specific activity in the regular assay in comparison with the *TDH3* promoter. In the regular assay a specific activity of 0.24-0.32 U/mg (Fig. 2 and 3) or 0.45-0.50 U/mg (fig. S7) is measured in the Δ*leu1* yeast strain with *LEU1-* or *TDH3*-driven Leu1 expression, respectively.

**Fig. S6.**
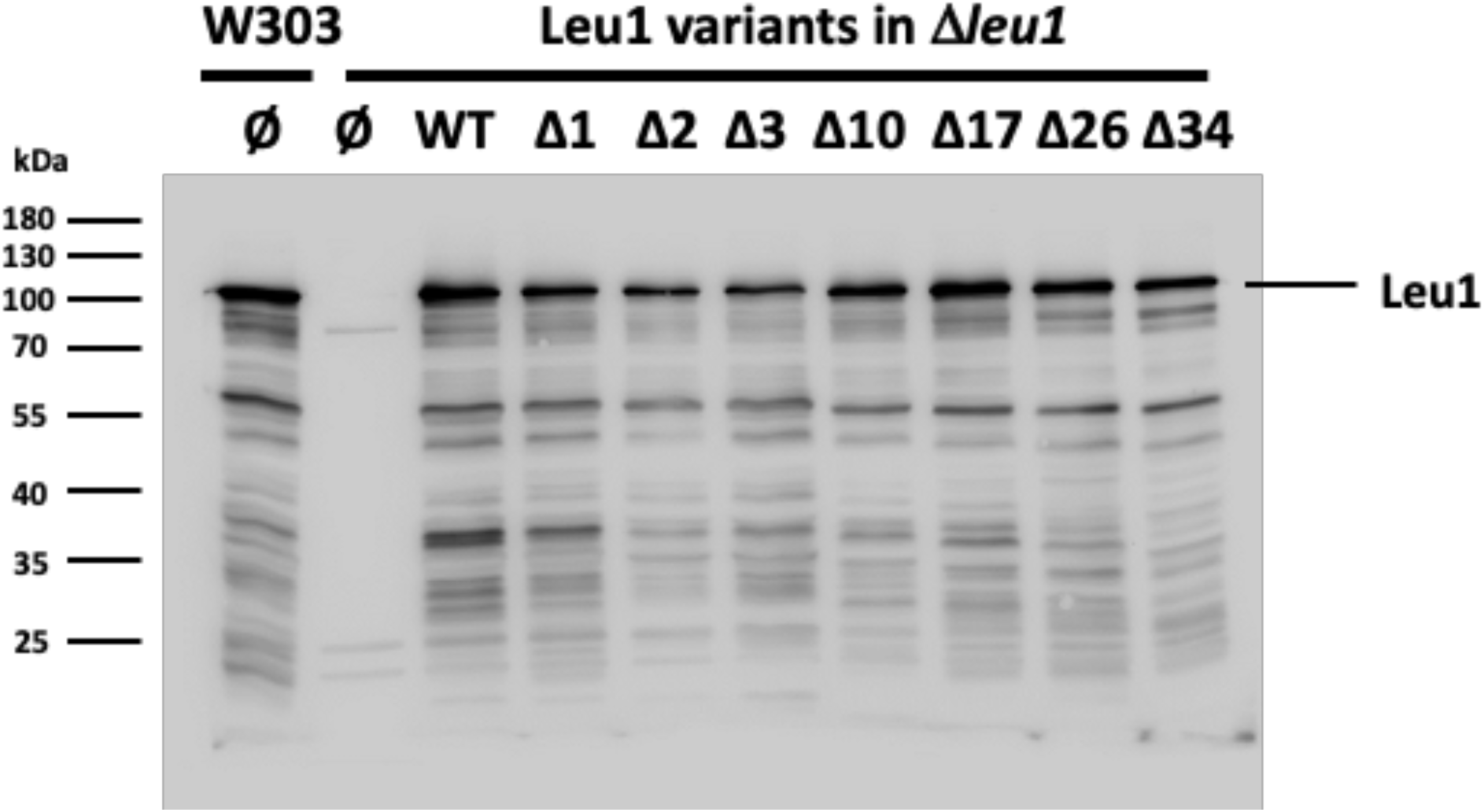
Western blot of cell free extracts of the indicated yeast strains transformed with pRS416 plasmids expressing wild type (WT) Leu1 and its C-terminally truncated variants from its natural promoter (corresponding to Fig. 2E). Rabbit polyclonal antibodies raised against purified yeast Leu1 were used.

**Fig. S7.**
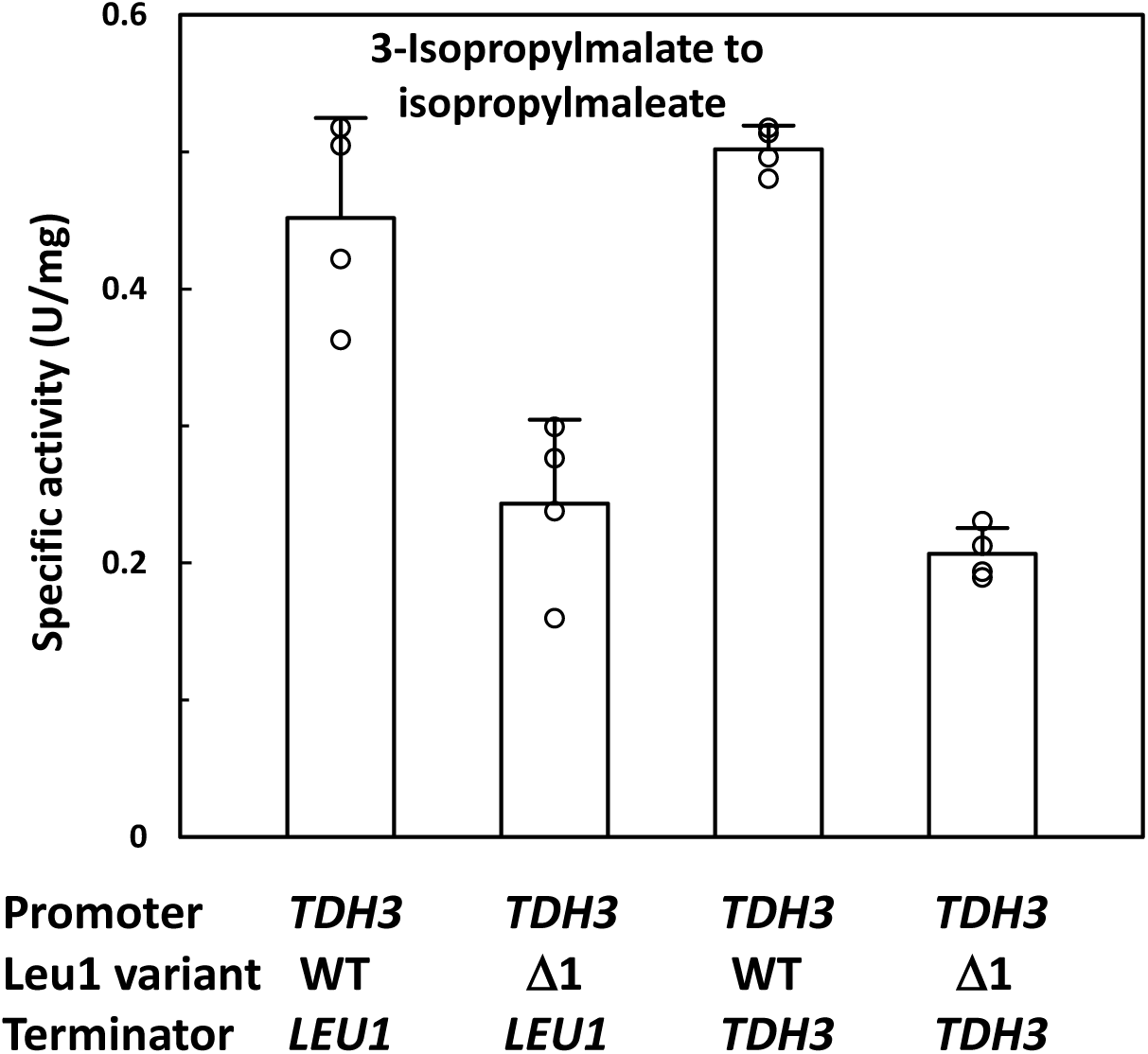
The loss of activity upon removal of W779 is not due to regulatory phenomena related to *LEU1* promoter or terminator regions. Specific activities (using the regular assay) of wild-type and the Δ1 truncation of yeast Leu1 isopropylmalate isomerase in cell extracts from Δ*leu1* cells transformed with pRS416 plasmids with *LEU1* under control of the indicated promoter or terminator.

**Fig. S8.**
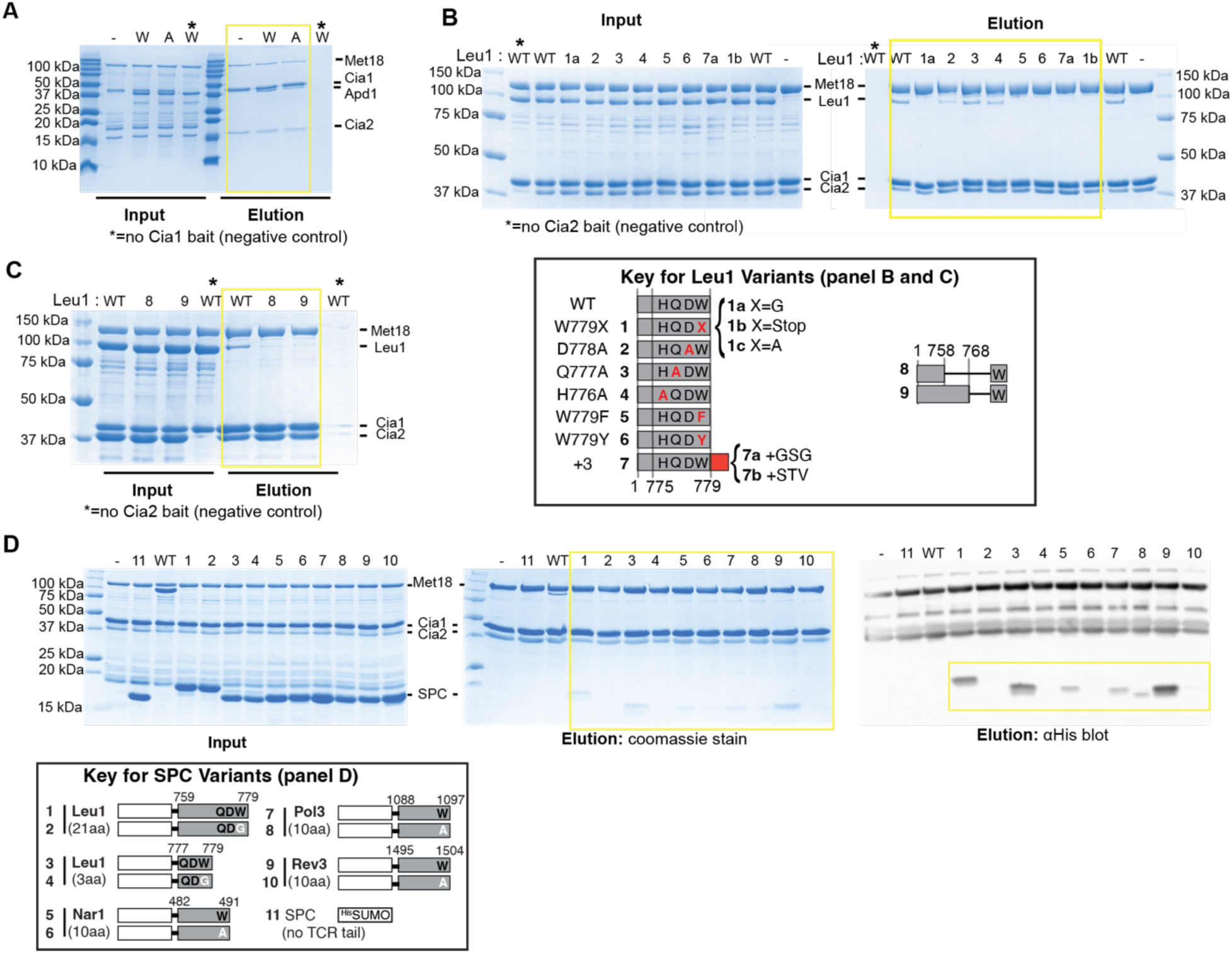
Expanded copurification data for Fig. 3A (**A**), Fig. 3C (**B**), Fig. 3F (**C**), and Fig. 4A (**D**). For each panel, SDS-PAGE gels (A-C) and Western Blot (C) from copurification input and elution samples are shown with data appearing in main text boxed in yellow. Numbers across the top of each gel indicate the Leu1 or SPC-variant (see keys included in figure) included in the copurification. In all panels, a positive control with the full-length, wild-type (WT) Leu1 or Apd1 protein is also shown. Panels A - C have a negative control (*) in which the ^Strep^Cia1 (A) or ^DT^Cia2 (B-C) bait was omitted. Additional controls in Panels A, B and D contain only the CTC, but no TCR-tail protein (−). The higher molecular weight bands in the Western blot correspond to the CTC subunits, which all have a His-tag to facilitate their purification.

**Fig. S9.**
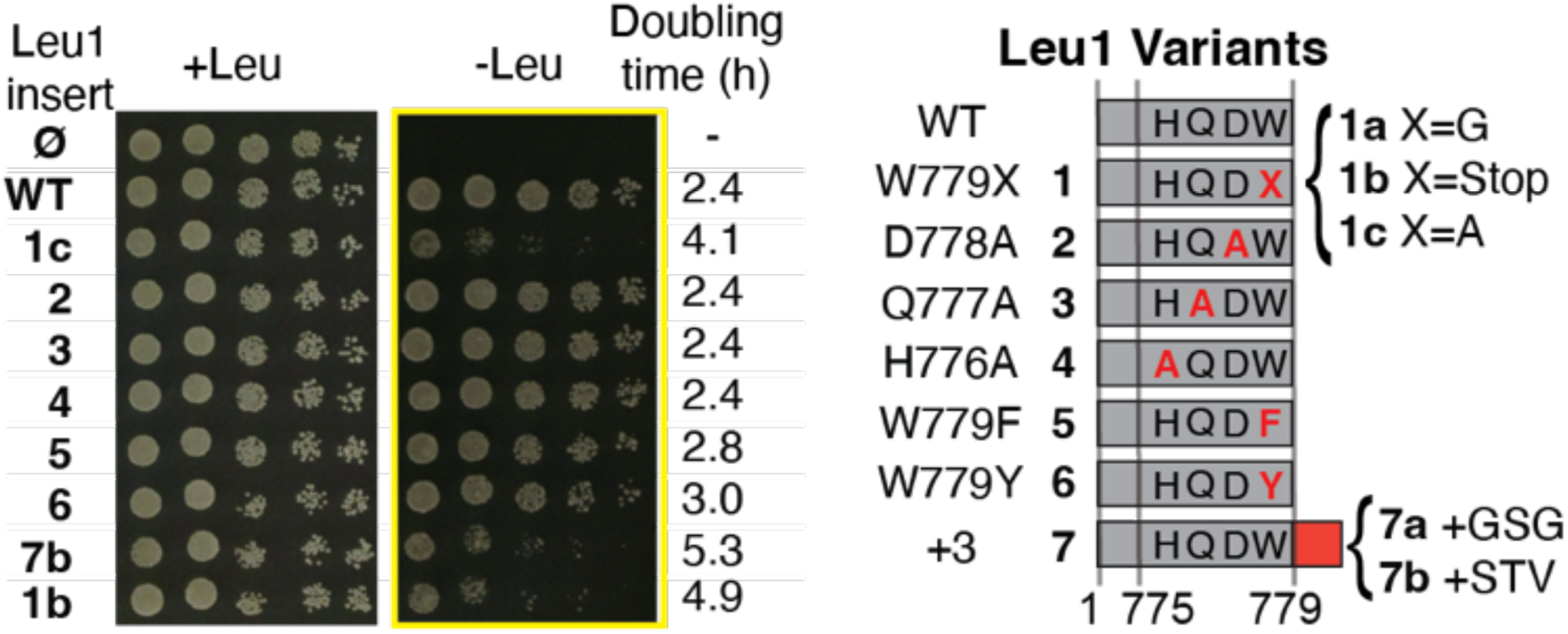
Full spot tests corresponding to Fig. 3D (yellow box). The left panel is a control plate onto which the same cell suspensions were spotted onto solid media supplemented with leucine. The doubling times measured in leucine deficient liquid medium are indicated on the right.

**Fig. S10.**
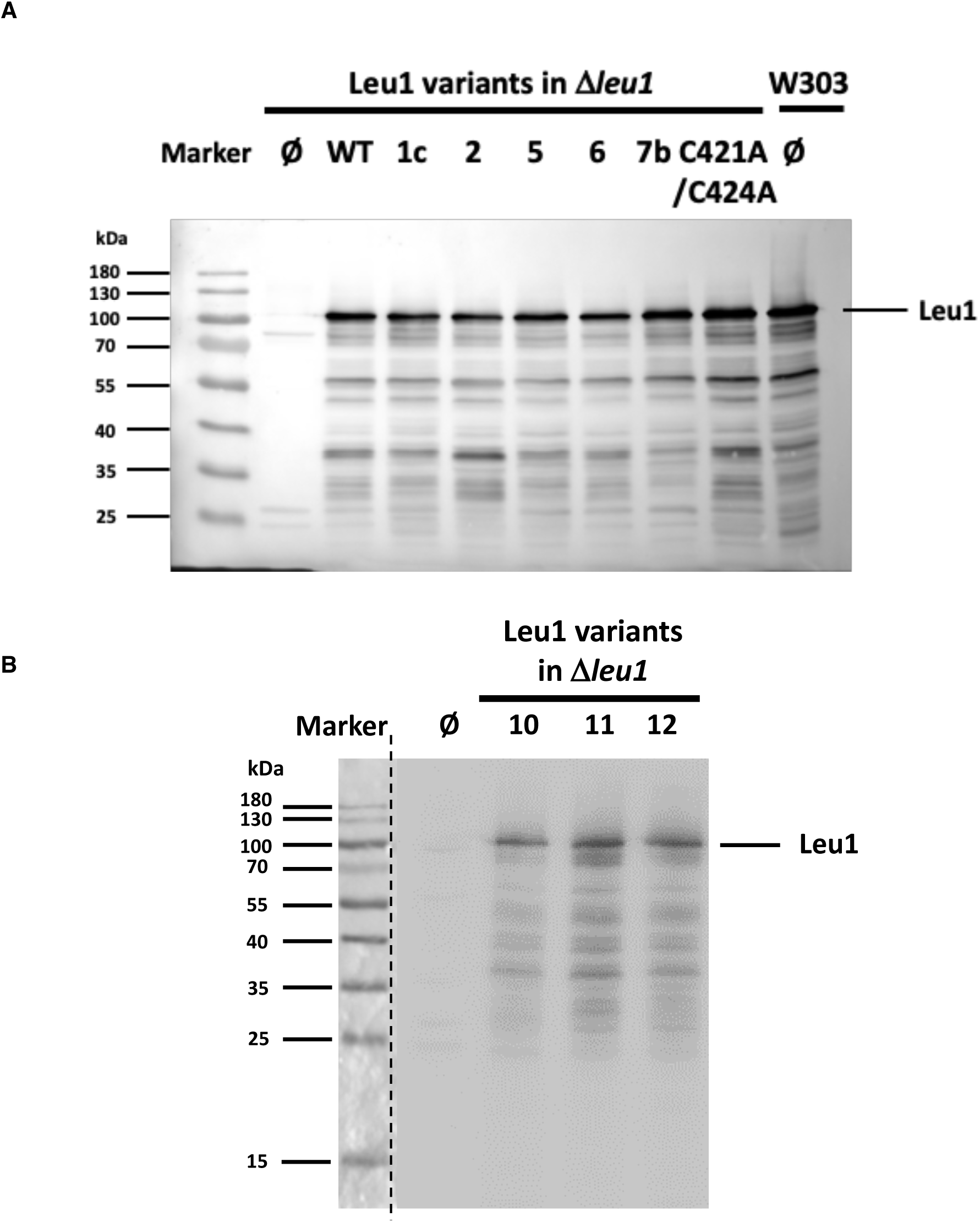
Western blots of cell free extracts of the indicated yeast strains transformed with 416 plasmids expressing wild type (WT) Leu1 and its variants from its natural promoter. Empty vector, Ø. (**A**) for Fig. 3E. (**B**) for Fig. 3G. Rabbit polyclonal antibodies raised against purified yeast Leu1 were used. The protein marker in (A) was merged on top of the chemiluminescence image by the Intas ChemoStar Touch imager software, for (B) the photograph (left of the dotted line) and chemiluminescence image were manually merged.

**Fig. S11.**
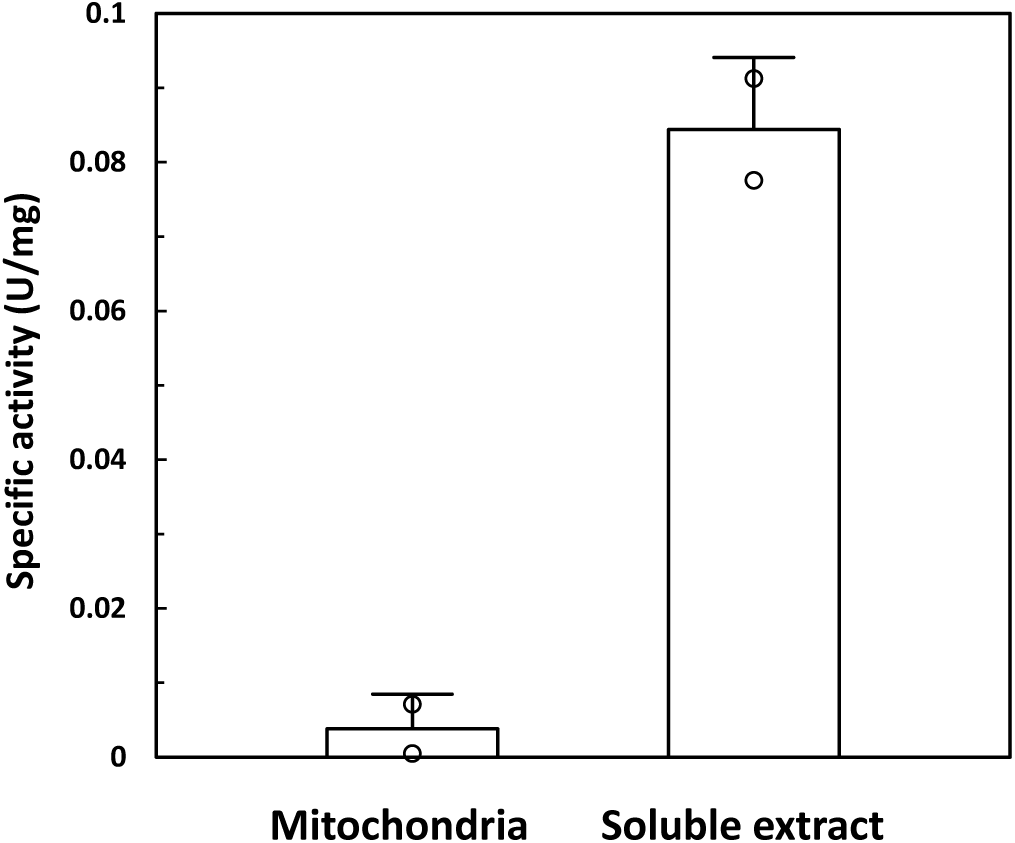
Fractionation of yeast expressing the engineered *E. coli* LeuCD with yeast C-terminal sequence demonstrates that the activity in the cell extract is cytosolic. The succinate dehydrogenase specific activity of the two mitochondrial preparations was 0.25 and 0.31 U/mg.

**Fig. S12.**
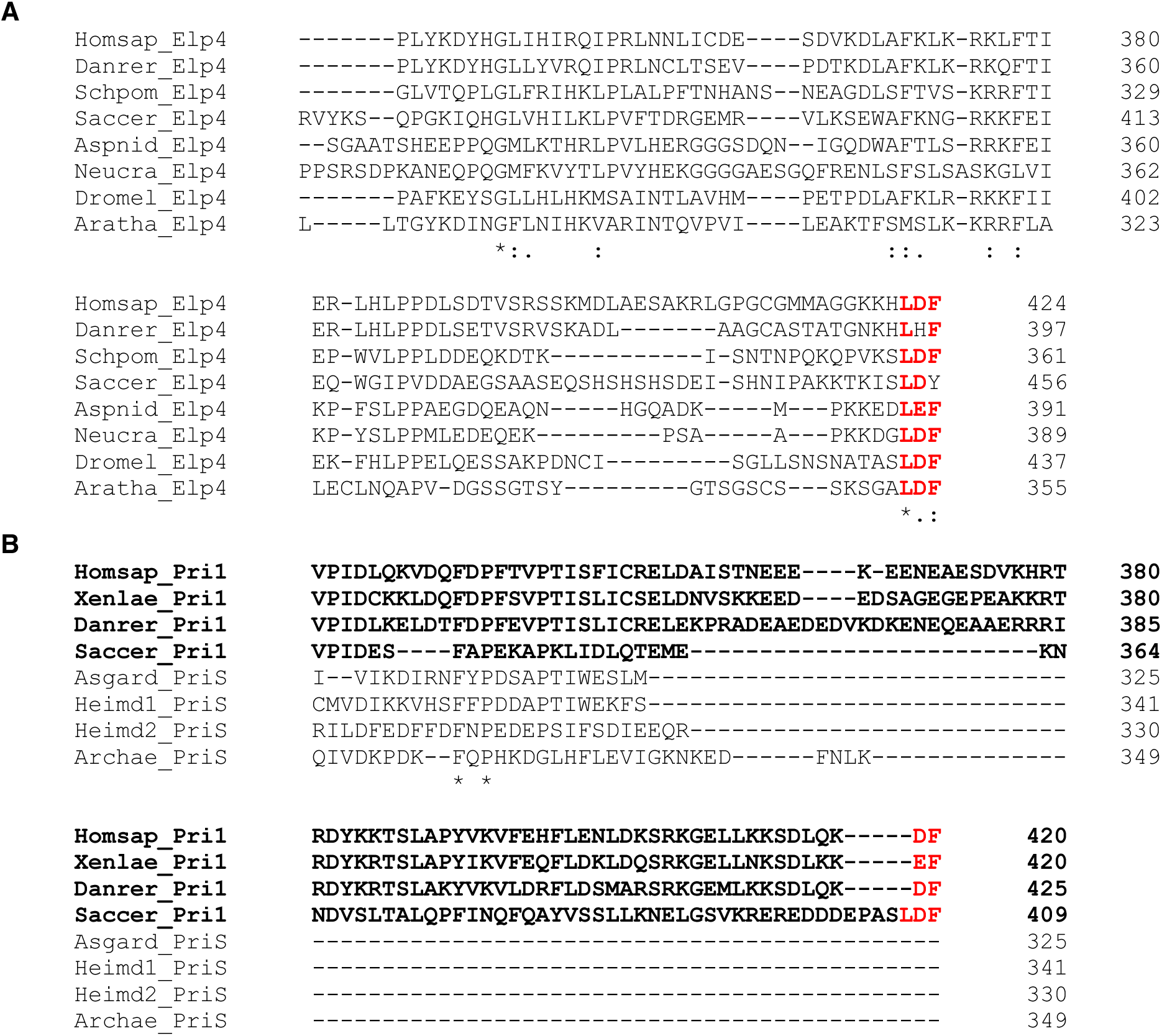
Amino acid sequence alignment for TCR-containing Elp4 and Pri1 non-Fe-S subunits of elongator and primase, respectively. (**A**) Elp4 subunits (Homsap, *Homo sapiens*; Danrer, *Danio rerio*; Schpom, *Schizosaccharomyces pombe*; Saccer, *Saccharomyces cerevisiae;* Aspnid, *Aspergillus nidulans*; Neucra, *Neurospora crassa*; Dromel, *Drosophila melanogaster*; Aratha, *Arabidopsis thaliana*) end in a TCR signal (red). Archaea and Eubacteria do not have Elongator complexes. The Elongator complex of *A. thaliana* is cytosolic. (**B**) Primase small subunits from Fungi and animals (bold) have a 40-60 amino acid extension and end in a TCR signal (red). In contrast, the primase small subunits from Asgard/Heimdall Archaeota, which are the closest non-eukaryotic homologs, do not have this extension. Homsap, *Homo sapiens*; Xenlae, *Xenopus laevis*; Danrer, *Danio rerio*; Saccer, *Saccharomyces cerevisiae*; Asgard_PriS, NPD89472.1; Heimd1_PriS, MBN1330204.1; Heimd2_PriS, PWI48945.1; Archae_PriS, NDB28160.1).

**Table S1.** Overview of failed expression of active Fe-S enzymes in heterologous hosts for biotechnological purposes.

| Pathway | Product | Reference |
| --- | --- | --- |
| Entner-Doudoroff | Ethanol | (47) |
| Entner-Doudoroff | Isobutanol | (46) |
| Mosaik | Isobutanol | (48) |
| Ehrlich | Isobutanol | (49) |
| Dahms (D-Xylose oxidation) | Ethyleneglycol | (50) |
| Weimberg | 2-Ketoglutarate | (51) |
| 2-C-Methyl-D-erythritol-4-phosphate (MEP) | Isoprenoids | (52) |
| 2-C-Methyl-D-erythritol-4-phosphate (MEP) | Isoprenoids | (53) |
| 1-Deoxy-D-xylulose-5-phosphate (DXP) | Isoprenoids | (54) |

**Table S2.** Proteomic data and structural data for TCR-containing proteins demonstrating that the TCR motif is not processed after Fe-S protein maturation.

| Source<sup>1</sup> | Protein Name | Occurrence<sup>2</sup> | Coverage (%)<sup>1</sup> | Calc. mass<sup>3</sup> | C-terminal peptide(s) observed and notes |
| --- | --- | --- | --- | --- | --- |
| *Homo sapiens* | POLD1<sup>4</sup> | - | - | - | Up to W1107 in structure |
|  | POLD1 | 170 | 98.9 | 956.4 | (K)(DLE)(DQEQLLR)(R)FGPPGPEAW |
|  | PUR1 | 72 | 100 | 934.4 | (E)(K)(SGH)(CTACL)(TG)(K)YPVELEW |
|  | NARF | 15 | 89.2 | 204.1 | (R)(G)THSLDIKW, non-tryptic digest |
|  | PRIM1 | 4 | 97.3 | 280.1 | (LLKK)SDLQKDF, non-tryptic and incomplete tryptic digests |
|  | ELP4 | 3 | 91.5 | 530.2 | (DLAE)SAKRLGPGCGMMAGGKKHLDF, non-tryptic digest |
|  | CIAO3 | 2 | 98.3 | 204.1 | ASTGLGIRW, incomplete tryptic digest |
|  | LTO1 | 2 | 95.6 | 966.5 | ISAEGSGLSF |
|  | POLR3F | 1 | 93.9 | 3561.5 | APCGLCPVFDDCHEGGEISPSNCIYMTEWLEF |
|  | CTU1 | 1 | 88.5 | 533.3 | PARPPASKAVPTF, non-tryptic digest |
|  | REV3L<sup>5</sup> | 0 | 52.5 | 762.4 | Too small, QLLDQF |
|  | RSAD2<sup>5</sup> | 0 | 62.0 | 432.2 | Too small, LDW |
| *Saccharomyces cerevisiae* | Leu1 | 27 | 100 | 683.3 | (K)AVTTTFDKVHQDW, incomplete tryptic digest |
|  | Pri1 | 21 | 89.9 | 1251.5 | (R)(ER)EDDDEPASLDF |
|  | Nar1 | 3 | 78.8 | 962.5 | DLVSVGSTW |
|  | Pol3<sup>6</sup> | - | - | - | Up to W1097 in structure |
|  | Pol3 | 1 | 81.7 | 204.1 | VEQLSKW, incomplete tryptic digest |
|  | Lto1<sup>7</sup> | 1 | 83.3 | 860.4 | QNQAQSW |
|  | Rev3<sup>5</sup> | 0 | 12.9 | 1188.6 | EEALISLNDW |
|  | Apd1<sup>5</sup> | 0 | 94.3 | 582.2 | Too small, YQSW |
|  | Ncs6<sup>5</sup> | 0 | 72.4 | 365.2 | Too small, LSF |
|  | Elp4<sup>5</sup> | 0 | 74.3 | 609.3 | Too small, ISLDY |

<sup>1</sup> Mass spectrometric data were collected with the PeptideAtlas online tool (55).
<sup>2</sup> The number of experiments for which the C-terminal peptide was detected in the canonical isoform.
<sup>3</sup> The calculated masses (monoisotopic) are for the C-terminal tryptic peptide.
<sup>4</sup> PolD1 was expressed in insect cells and its structure was determined by cryoEM (PDB 61SM).
<sup>5</sup> Failure to detect C-terminal peptides explained by the small size of the C-terminal peptide (<7 amino acids) or the moderate coverage due to the low cellular protein abundance.
<sup>6</sup> Pol3 was expressed in yeast and its structure was determined by X-ray crystallography (PDB 6P1H)
<sup>7</sup> Lto1 coverage corrected to take the assignment of the N-terminal sequence (MDF) into account which is supported by the absence of peptides corresponding to the erroneously assigned MVR N-terminus (22).

**Table S3.**
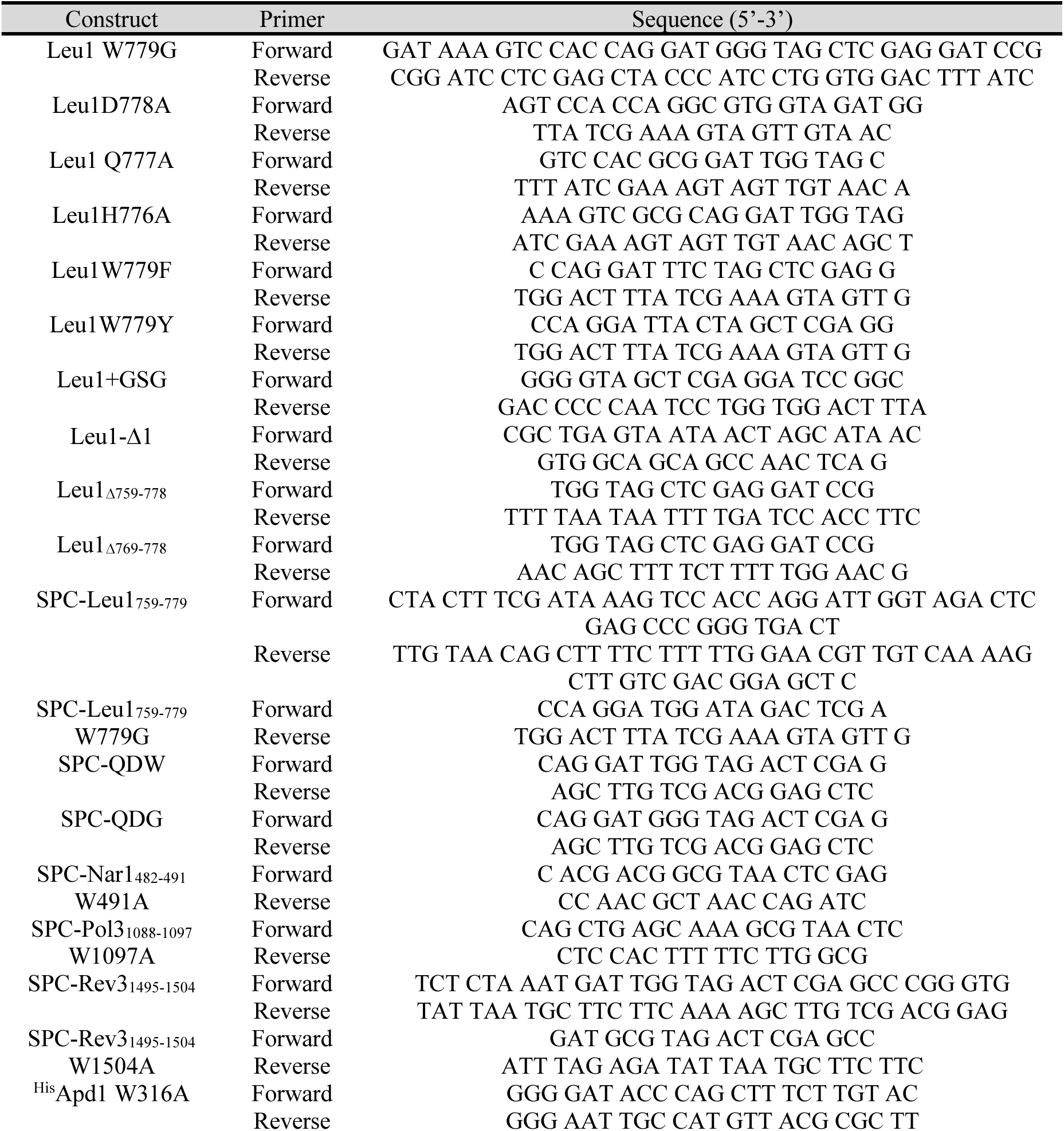
Mutagenesis primers for plasmids relating to *in vitro* interaction studies.

**Table S4.**
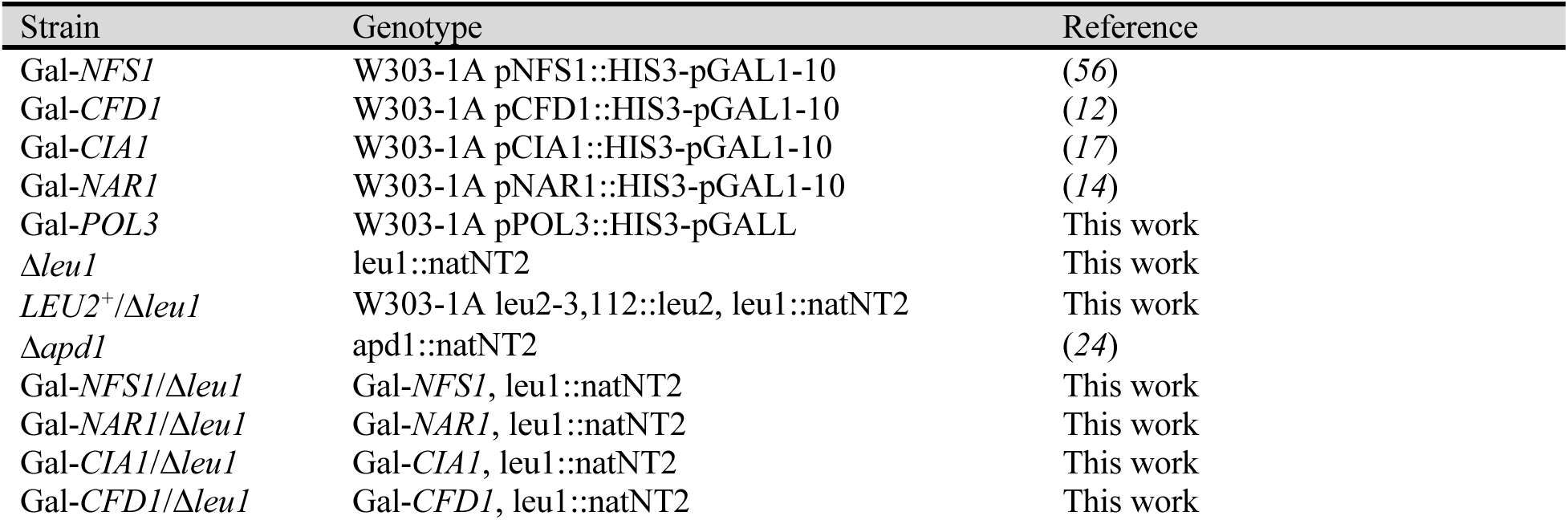
Yeast strains.

| Strain | Genotype | Reference |
| --- | --- | --- |
| Gal-*NFS1* | W303-1A pNFS1::HIS3-pGAL1-10 | (56) |
| Gal-*CFD1* | W303-1A pCFD1::HIS3-pGAL1-10 | (12) |
| Gal-*CIA1* | W303-1A pCIA1::HIS3-pGAL1-10 | (17) |
| Gal-*NAR1* | W303-1A pNAR1::HIS3-pGAL1-10 | (14) |
| Gal-*POL3* | W303-1A pPOL3::HIS3-pGALL | This work |
| $\Delta leu1$ | leu1::natNT2 | This work |
| *LEU2*<sup>+</sup>/$\Delta leu1$ | W303-1A leu2-3,112::leu2, leu1::natNT2 | This work |
| $\Delta apd1$ | apd1::natNT2 | (24) |
| Gal-*NFS1*/$\Delta leu1$ | Gal-*NFS1*, leu1::natNT2 | This work |
| Gal-*NAR1*/$\Delta leu1$ | Gal-*NAR1*, leu1::natNT2 | This work |
| Gal-*CIA1*/$\Delta leu1$ | Gal-*CIA1*, leu1::natNT2 | This work |
| Gal-*CFD1*/$\Delta leu1$ | Gal-*CFD1*, leu1::natNT2 | This work |

**Table S5:**
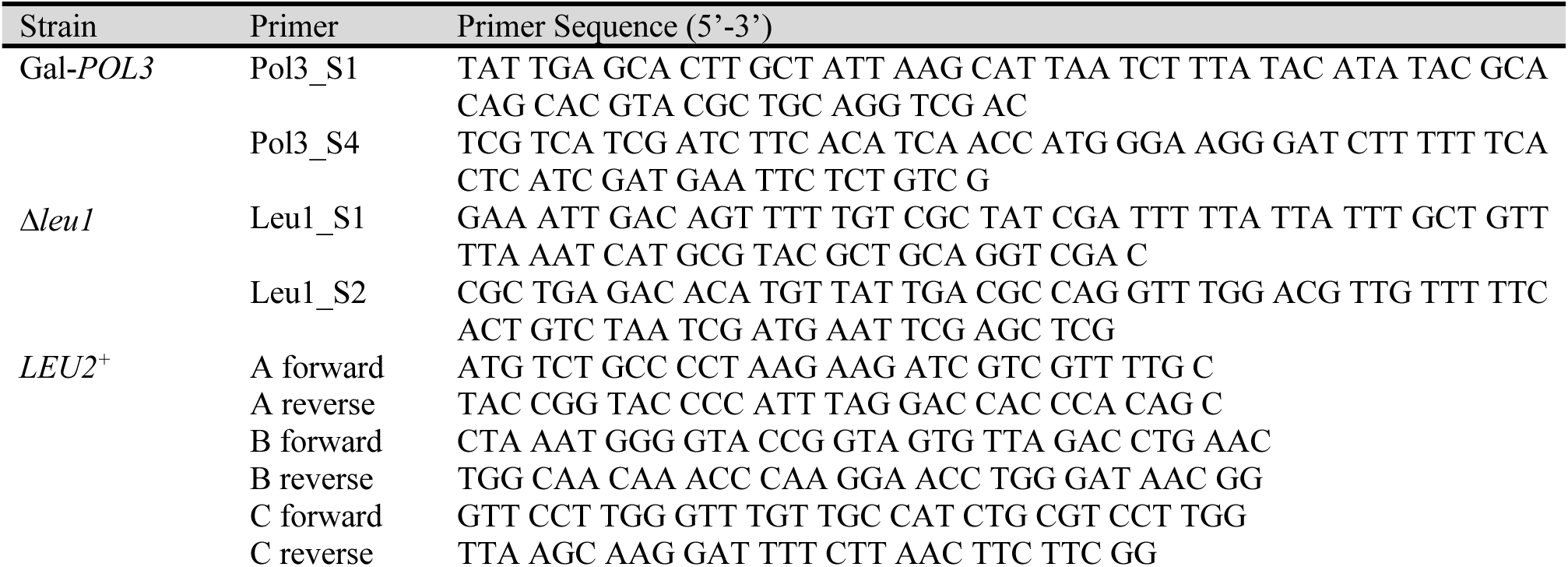
Primers for homologous recombination.

**Table S6.**
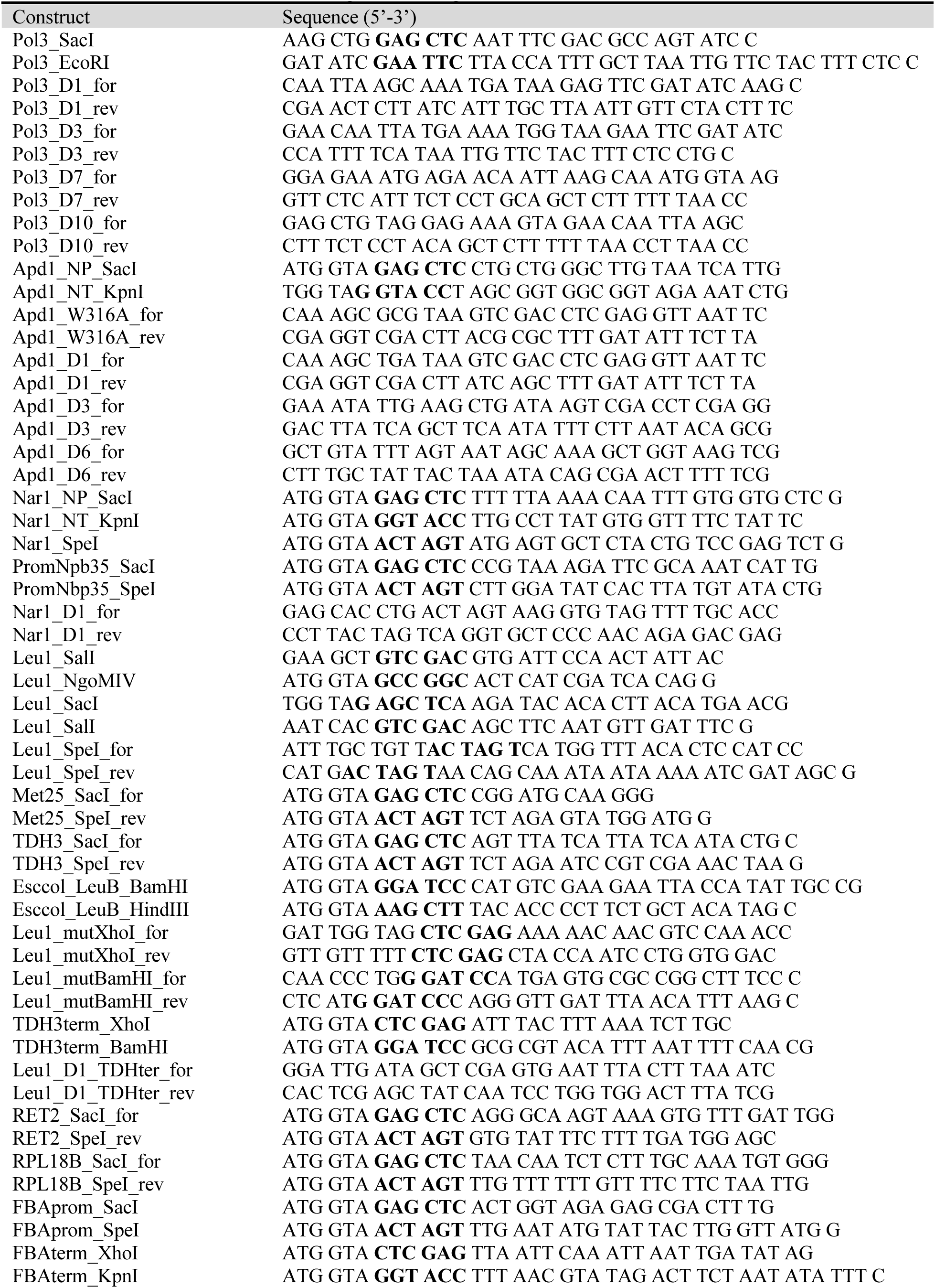
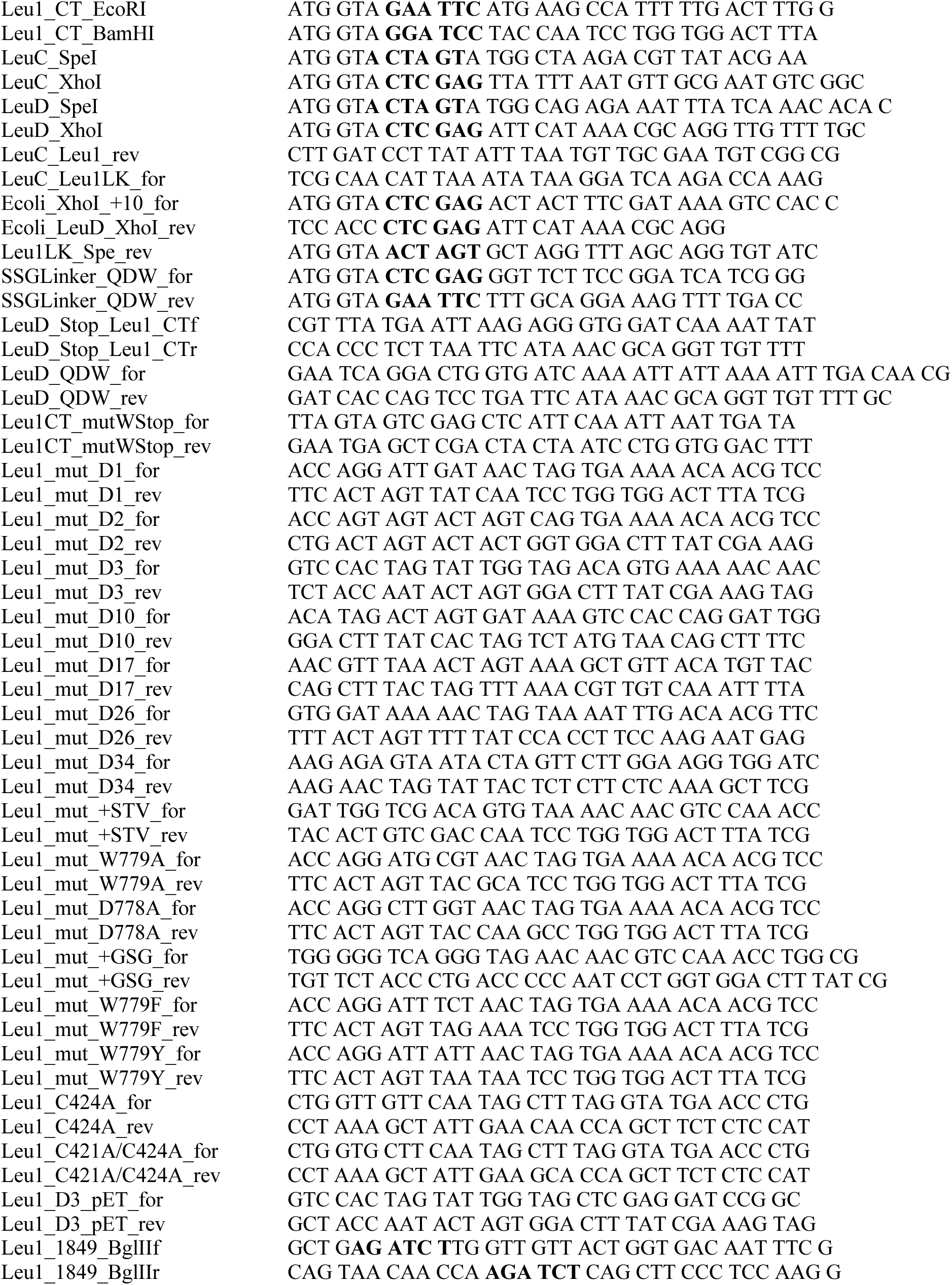
Primers used for cloning and mutagenesis. Restriction sites are in bold.

**Table S7:**
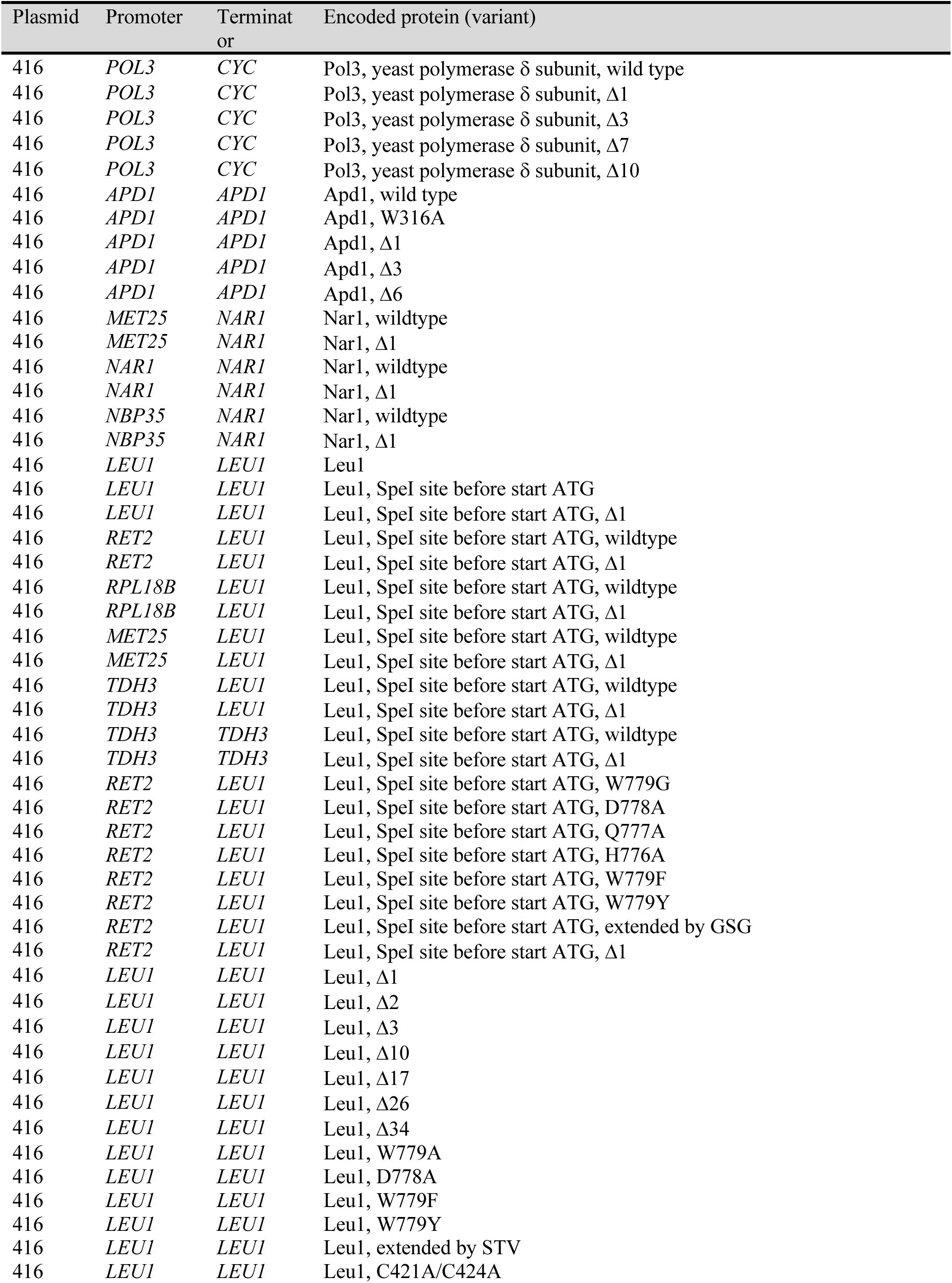
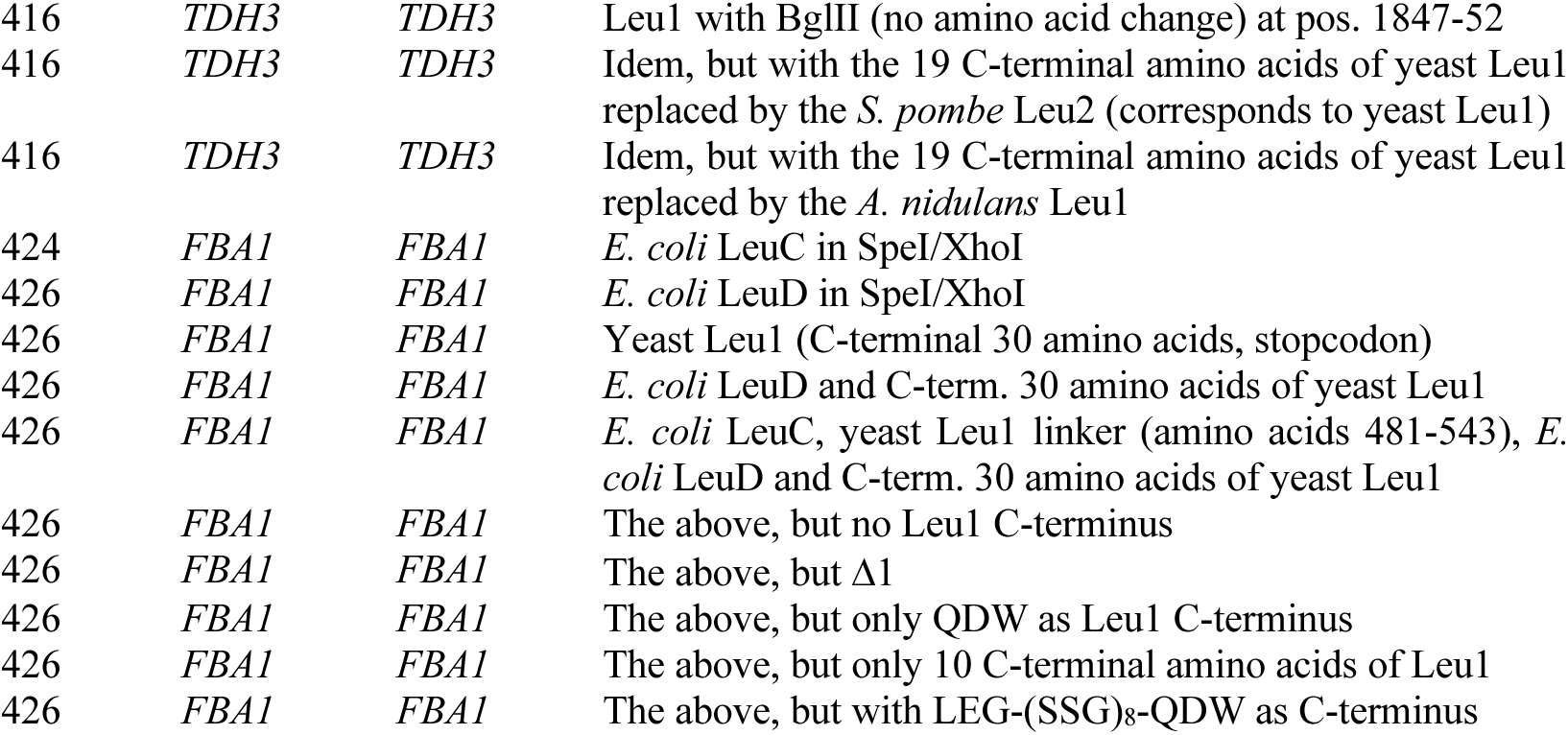
Yeast plasmids. Δ refers to C-terminal truncation.

| Plasmid | Promoter | Terminat<br>or | Encoded protein (variant) |
| --- | --- | --- | --- |
| 416 | *POL3* | *CYC* | Pol3, yeast polymerase $\delta$ subunit, wild type |
| 416 | *POL3* | *CYC* | Pol3, yeast polymerase $\delta$ subunit, $\Delta 1$ |
| 416 | *POL3* | *CYC* | Pol3, yeast polymerase $\delta$ subunit, $\Delta 3$ |
| 416 | *POL3* | *CYC* | Pol3, yeast polymerase $\delta$ subunit, $\Delta 7$ |
| 416 | *POL3* | *CYC* | Pol3, yeast polymerase $\delta$ subunit, $\Delta 10$ |
| 416 | *APD1* | *APD1* | Apd1, wild type |
| 416 | *APD1* | *APD1* | Apd1, W316A |
| 416 | *APD1* | *APD1* | Apd1, $\Delta 1$ |
| 416 | *APD1* | *APD1* | Apd1, $\Delta 3$ |
| 416 | *APD1* | *APD1* | Apd1, $\Delta 6$ |
| 416 | *MET25* | *NAR1* | Nar1, wildtype |
| 416 | *MET25* | *NAR1* | Nar1, $\Delta 1$ |
| 416 | *NAR1* | *NAR1* | Nar1, wildtype |
| 416 | *NAR1* | *NAR1* | Nar1, $\Delta 1$ |
| 416 | *NBP35* | *NAR1* | Nar1, wildtype |
| 416 | *NBP35* | *NAR1* | Nar1, $\Delta 1$ |
| 416 | *LEU1* | *LEU1* | Leu1 |
| 416 | *LEU1* | *LEU1* | Leu1, SpeI site before start ATG |
| 416 | *LEU1* | *LEU1* | Leu1, SpeI site before start ATG, $\Delta 1$ |
| 416 | *RET2* | *LEU1* | Leu1, SpeI site before start ATG, wildtype |
| 416 | *RET2* | *LEU1* | Leu1, SpeI site before start ATG, $\Delta 1$ |
| 416 | *RPL18B* | *LEU1* | Leu1, SpeI site before start ATG, wildtype |
| 416 | *RPL18B* | *LEU1* | Leu1, SpeI site before start ATG, $\Delta 1$ |
| 416 | *MET25* | *LEU1* | Leu1, SpeI site before start ATG, wildtype |
| 416 | *MET25* | *LEU1* | Leu1, SpeI site before start ATG, $\Delta 1$ |
| 416 | *TDH3* | *LEU1* | Leu1, SpeI site before start ATG, wildtype |
| 416 | *TDH3* | *LEU1* | Leu1, SpeI site before start ATG, $\Delta 1$ |
| 416 | *TDH3* | *TDH3* | Leu1, SpeI site before start ATG, wildtype |
| 416 | *TDH3* | *TDH3* | Leu1, SpeI site before start ATG, $\Delta 1$ |
| 416 | *RET2* | *LEU1* | Leu1, SpeI site before start ATG, W779G |
| 416 | *RET2* | *LEU1* | Leu1, SpeI site before start ATG, D778A |
| 416 | *RET2* | *LEU1* | Leu1, SpeI site before start ATG, Q777A |
| 416 | *RET2* | *LEU1* | Leu1, SpeI site before start ATG, H776A |
| 416 | *RET2* | *LEU1* | Leu1, SpeI site before start ATG, W779F |
| 416 | *RET2* | *LEU1* | Leu1, SpeI site before start ATG, W779Y |
| 416 | *RET2* | *LEU1* | Leu1, SpeI site before start ATG, extended by GSG |
| 416 | *RET2* | *LEU1* | Leu1, SpeI site before start ATG, $\Delta 1$ |
| 416 | *LEU1* | *LEU1* | Leu1, $\Delta 1$ |
| 416 | *LEU1* | *LEU1* | Leu1, $\Delta 2$ |
| 416 | *LEU1* | *LEU1* | Leu1, $\Delta 3$ |
| 416 | *LEU1* | *LEU1* | Leu1, $\Delta 10$ |
| 416 | *LEU1* | *LEU1* | Leu1, $\Delta 17$ |
| 416 | *LEU1* | *LEU1* | Leu1, $\Delta 26$ |
| 416 | *LEU1* | *LEU1* | Leu1, $\Delta 34$ |
| 416 | *LEU1* | *LEU1* | Leu1, W779A |
| 416 | *LEU1* | *LEU1* | Leu1, D778A |
| 416 | *LEU1* | *LEU1* | Leu1, W779F |
| 416 | *LEU1* | *LEU1* | Leu1, W779Y |
| 416 | *LEU1* | *LEU1* | Leu1, extended by STV |
| 416 | *LEU1* | *LEU1* | Leu1, C421A/C424A |
| 416 | *TDH3* | *TDH3* | Leu1 with BglII (no amino acid change) at pos. 1847-52 |
| 416 | *TDH3* | *TDH3* | Idem, but with the 19 C-terminal amino acids of yeast Leu1 replaced by the *S. pombe* Leu2 (corresponds to yeast Leu1) |
| 416 | *TDH3* | *TDH3* | Idem, but with the 19 C-terminal amino acids of yeast Leu1 replaced by the *A. nidulans* Leu1 |
| 424 | *FBA1* | *FBA1* | *E. coli* LeuC in SpeI/XhoI |
| 426 | *FBA1* | *FBA1* | *E. coli* LeuD in SpeI/XhoI |
| 426 | *FBA1* | *FBA1* | Yeast Leu1 (C-terminal 30 amino acids, stopcodon) |
| 426 | *FBA1* | *FBA1* | *E. coli* LeuD and C-term. 30 amino acids of yeast Leu1 |
| 426 | *FBA1* | *FBA1* | *E. coli* LeuC, yeast Leu1 linker (amino acids 481-543), *E. coli* LeuD and C-term. 30 amino acids of yeast Leu1 |
| 426 | *FBA1* | *FBA1* | The above, but no Leu1 C-terminus |
| 426 | *FBA1* | *FBA1* | The above, but $\Delta 1$ |
| 426 | *FBA1* | *FBA1* | The above, but only QDW as Leu1 C-terminus |
| 426 | *FBA1* | *FBA1* | The above, but only 10 C-terminal amino acids of Leu1 |
| 426 | *FBA1* | *FBA1* | The above, but with LEG-(SSG)<sub>8</sub>-QDW as C-terminus |

**Table S8:** *E. coli* expression plasmids. The wild type Leu1 plasmid was described in Netz et al.(*12*)

| Plasmid | Encoded protein (variant) |
| --- | --- |
| pET15b | N-terminally His-tagged Leu1, wild type |
| pET15b | N-terminally His-tagged Leu1, $\Delta 3$ |
| pET15b | N-terminally His-tagged Leu1, $\Delta 10$ |
| pET15b | N-terminally His-tagged Leu1, $\Delta 17$ |
| pET15b | N-terminally His-tagged Leu1, $\Delta 34$ |
| pET-Duet1 | N-terminally His-tagged *E. coli* LeuB |

**Data S1.**

Tabulation of C-terminal proteomes of Fe-S proteins in *H. sapiens, S. cerevisiae, A thaliana, E. coli,* and *M. jannaschii*.

**Data S2.**

Analysis of C-terminal tails of CIA clients, factors, and adaptors from 10 eukaryotic reference organisms.

